# High-throughput analysis of dendritic and axonal arbors reveals transcriptomic correlates of neuroanatomy

**DOI:** 10.1101/2022.03.07.482900

**Authors:** Olga Gliko, Matt Mallory, Rachel Dalley, Rohan Gala, James Gornet, Hongkui Zeng, Staci Sorensen, Uygar Sümbül

**Affiliations:** Allen Institute, Seattle WA, USA; California Institute of Technology, Pasadena, CA, USA

## Abstract

Neuronal anatomy is central to the organization and function of brain cell types. However, anatomical variability within apparently homogeneous populations of cells can obscure such insights. Here, we report large-scale automation of neuronal morphology reconstruction and analysis on a dataset of 813 inhibitory neurons characterized using the Patch-seq method, which enables measurement of multiple properties from individual neurons, including local morphology and transcriptional signature. We demonstrate that these automated reconstructions can be used in the same manner as manual reconstructions to understand the relationship between some, but not all, cellular properties used to define cell types. We uncover gene expression correlates of laminar innervation on multiple transcriptomically defined neuronal subclasses and types. In particular, our results reveal correlates of the variability in Layer 1 (L1) axonal innervation in a transcriptomically defined subpopulation of Martinotti cells in the adult mouse neocortex.

## Introduction

The shape of dendrites and axons, their distribution within the neuropil, and patterns of their long-range projections can reveal fundamental principles of nervous system organization and function. In the cortex, much of our understanding depends on the anatomical and functional descriptions of cortical layers. Yet, the origin and role of morphological and molecular diversity of individual neurons within cortical layers beyond broad subclass identities is poorly understood, in part due to low sample numbers. While molecular profiling techniques have recently improved by orders of magnitude, anatomical characterization remains time consuming due to continued reliance on (semi-)manual reconstruction.

Improvements in the throughput of the Patch-seq technique^1–8^ have enabled measurement of electrophysiological features, transcriptomic signatures, and local morphology in slice prepa-rations for thousands of neurons in recent studies.^5, 6^ In these repetitive experiments where maintaining a high throughput is a primary goal,^9^ the brightfield microscope’s speed, ease of use, and ubiquity make it an attractive choice to image local neuronal morphology. While this choice helps to streamline the experimental steps, morphological reconstruction remains a major bottleneck of overall throughput, in part due to limited imaging resolution, even with state-of-the-art semi-manual tools.^5^

A rich literature exists on automated segmentation in sparse imaging scenarios. However, these methods typically focus on high-contrast, high-resolution images obtained by optical sectioning microscopy (i.e., confocal, two-photon, and light-sheet),^10–18^ and are not immediately applicable to brightfield images because of the significantly worse depth resolution and the complicated point spread function of the brightfield microscope. Moreover, segmentation of full local morphology together with identification of the axon, dendrites, and soma has remained elusive for methods tested on image stacks obtained by the brightfield microscope.^19–22^ Therefore, we first introduce an end-to-end automated neuron reconstruction pipeline (Figure 1a) to improve scalability of brightfield 3D image-based reconstructions in Patch-seq experiments by a few orders of magnitude. We note that our primary goal is not to report on a more accurate reconstruction method *per se*. Rather, we aim to demonstrate how automated tracing, with its potential mistakes, can be leveraged to rigorously address certain scientific questions by increasing the throughput. To this end, we select a set of brightfield images and use the corresponding manually reconstructed neuron traces to assign voxel-wise ground truth labels (axon, dendrite, soma, background). Next, we design a custom deep learning model and train it on this curated ground truth dataset to perform 3D image segmentation using volumetric patches of the raw image as input. We implement fully automated post-processing steps, including basal vs. apical dendrite separation for pyramidal cells, for the resulting segmentations to obtain annotated traces for each neuron. We compare the accuracy of these automated traces with a held out set of manual reconstructions, based on geometrical precision, a suite of morphometric features, and arbor density representations derived from the traces.

We utilize this pipeline to reconstruct a large set of neurons from Patch-seq experiments, and use the transcriptomic profiles captured from the same cells to systematically search for gene subsets that can predict certain aspects of neuronal anatomy. The existence of a hierarchical transcriptomic taxonomy^23^ enables studying subsets of neurons at different levels of the transcriptomic hierarchy. At the finest scale of the hierarchy (transcriptomic types or “t-types”^5^), we study seven interneuron types and focus on a transcriptomically defined sub-population of L1-projecting, *Sst* gene expressing neurons (Sst cells) that correspond to Martinotti cells (See, for instance, Ref.^24^). While previous studies have elucidated the role of Martinotti cells in gating top-down input to pyramidal neurons via their L1-innervating axons,^25, 26^ the wide variability in the extent of L1 innervation behind it is not well understood. Our results suggest transcriptomic correlates of the innervating axonal mass, which may control the amount of top-down input to canonical cortical circuits. Our approach represents a general program to systematically connect gene expression with neuronal anatomy in a high-throughput and data-driven manner.

## Results

### An automated morphology reconstruction pipeline for brightfield microscopy

As the first step to automate the reconstruction of in-slice brightfield images of biocytin-filled neurons, we curate a set of manually traced neurons. While this set should ideally be representative of the underlying image space, it should also be as small as possible to facilitate downstream cross-validation studies via an abundance of cells not used during training. We thus choose 51 manually traced neurons as the training set to represent the underlying variability in the image quality and neuronal identity (via the available t-type labels). We develop a topology-preserving variant of the fast marching algorithm^13^ to generate volumetric labels from manual traces (Figure 1b). We train a convolutional neural network (U-Net)^16, 27–29^ using image stacks and labels as the training set and employing standard data augmentation strategies to produce initial segmentations of neuronal morphologies (Figure 1b,c). While knowledge of axonal vs. dendritic branches informs most existing insight, their automated identification poses a challenge due to the limited field-of-view of artificial neural networks. We find that image and trace context that is in the vicinity of the initial segmentation is sufficient to correct many axon vs. dendrite labeling mistakes in an efficient way because this effectively reduces the problem to a single dimension, i.e., features calculated along the 1D initial trace (Figure 1d). We further algorithmically post-process the segmentations to correct connectivity mistakes introduced by the neural network and obtain the final reconstruction of axons and dendrites (Figure 1d, Figure 2a, Figures S1-S11, Methods). We observe that this approach offers marked improvements in tracing quality compared to a previous large-scale effort focusing on fluorescent, optical-sectioning microscopy (Figure S12). Moreover, Figure S13 shows that its segmentation quality remains robust when tested, without any tuning, with images from different species and brain structures.^30–32^ (Fine-tuning the existing model with a small training set representing the tissue of interest should further improve performance.) The overall pipeline produces neuron reconstructions in the commonly-used swc format^33^ from raw image stacks at a rate of *∼*6 cells/day with a single GPU card (Methods). Our setup uses 16 cards to achieve two orders of magnitude improvement in speed over semi-manual segmentation^5^ with one anatomist. We have so far processed the cells reported in Ref.,^5^ which mapped neurons to an existing taxonomy of transcriptomic cell types^23^ and introduced a transcription-centric, multimodal analysis of inhibitory neurons. We have also processed a set of *∼*700 excitatory neurons which are analyzed in Ref.^34^ (While we typically display only the apical and basal dendrite segmentations for excitatory cells, the method can also trace and label the local axon when it is captured in the slice (Figure S14).) After quality control steps (Methods), we focus on a set of 813 interneurons for further study in this paper.

**Figure 1:**
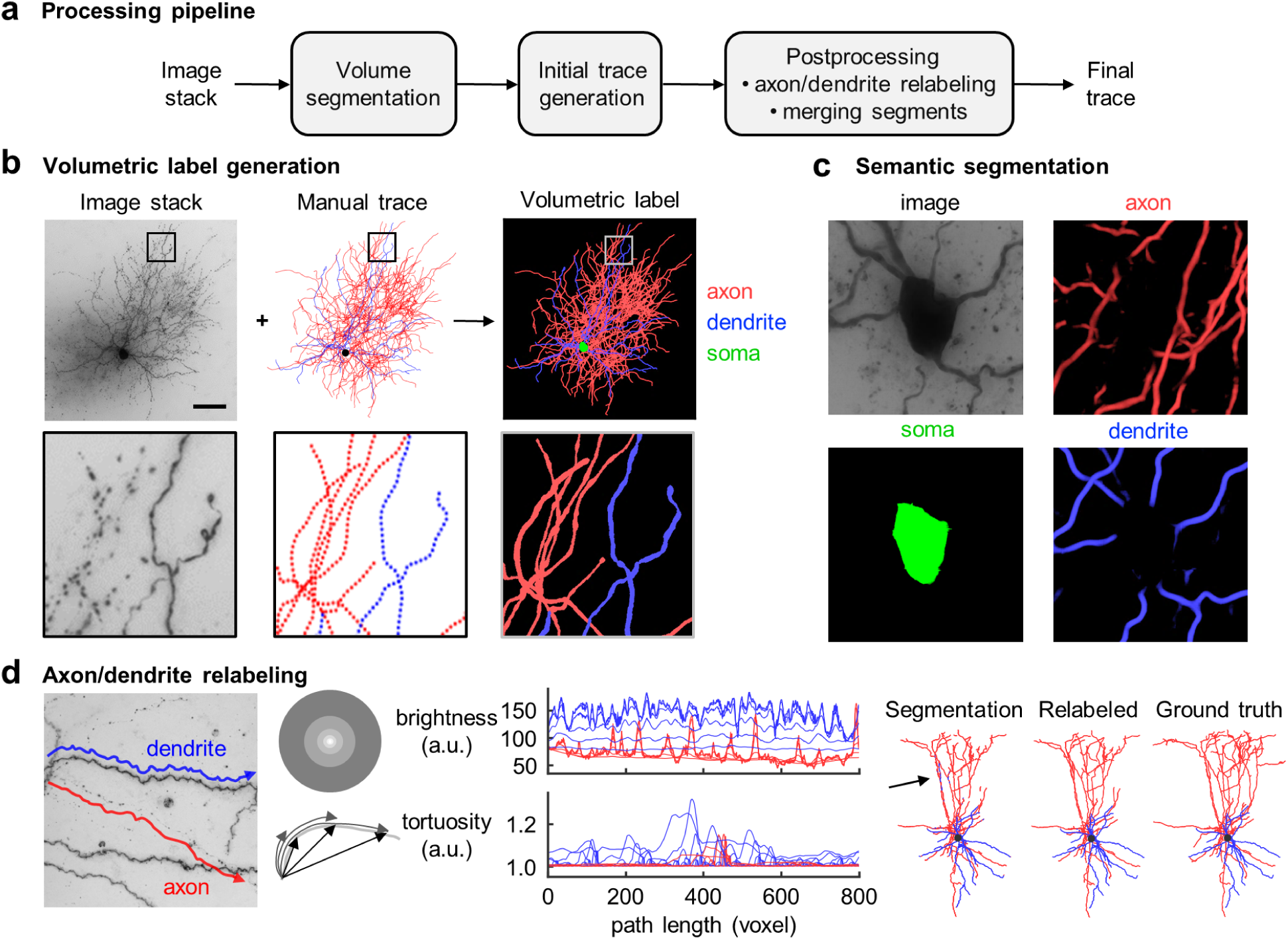
Neuron reconstruction pipeline for in-slice bright-field images of biocytin-filled neurons. **a**, Processing pipeline. Convolutional neural network (CNN) segmentations of 3D image stacks are post-processed by custom machine learning tools to produce digital representations of neuronal morphologies. **b**, Topology preserving fast marching algorithm generates the volumetric label from raw image stack and manual skeletonization. Dendrites (blue), axons (red), soma (green) are separately labeled to train a supervised CNN model. Scale bar, 100 *µm*. **c**, Semantic segmentation provides accurate soma location and boundary. **d**, Axon/dendrite relabeling. A neural network model predicts node labels from multiple image brightness and trace tortuosity features based on local contexts of different size along the initial trace. (left, example image of dendrite and axon segments; middle, corresponding feature plots; right, automated traces of test neuron with/without relabeling vs. manual trace). Arrow indicates nodes mislabeled by segmentation and corrected during post-processing.

The proposed pipeline produces end-to-end automated reconstructions in single-cell imaging scenarios. However, in practice, neurons are patched near each other to increase the throughput of physiological and transcriptomic characterization. The resulting image, which typically centers on the neuron of interest, can therefore contain neurites from other neurons. Neurites from off-target neurons within the image stack cannot be properly characterized because they rarely remain in the field of view. As part of algorithmic post-processing and quality control, disconnected segments are removed automatically when they remain relatively far from the cell of interest (Methods). When multiple neurons are patched in close proximity, quality control by a human is needed to check for and remove nearby extraneous branches. To ensure the integrity of presented results with minimal manual effort, the cell is not used if quality control suggests the existence of nearby branches and a manual trace is not available. If the manual trace already exists, we simulate manual branch removal based on a mask obtained from the manual trace (Methods). We report quantification results separately for reconstructions obtained with/without nearby branch removal.

### Evaluation of reconstruction accuracy

We evaluate the quality of automated traces by comparing them to the manual traces which we regard as the ground truth. To compare a pair of automated and manual traces, we perform a bi-directional nearest-neighbor search to find correspondence nodes in both traces within a certain distance.^13^ A node in the automated trace that has (does not have) a corresponding node in the manual trace is referred to as a true (false) positive node, and a node in the manual trace that does not have a corresponding node in the automated trace is referred to as a false negative node. We calculate this metric separately for axonal and dendritic nodes, as well as for all nodes regardless of the type, and compute corresponding precision, recall, and f1-score (harmonic mean of the precision and recall) values. These metrics indicate how well the automated trace captures the layout of the axon/dendrites/neurites in a reconstructed neuron. Figure 2b and Table S1 display that, at a search radius of 10*µm*, the mean f1-score is above 0.8 for both axonal and dendritic morphologies. Therefore, we expect these automatically generated traces to perform comparably to their manually generated counterparts in analyses that do not require a resolution better than 10*µm*, as we demonstrate below. (Figure S15 and Table S2 provide basic estimates of cross-human tracing discrepancy based on one test cell.)

To further assess the similarity in the arbor layout and other aspects of morphology that are not captured by the node correspondence study, we use standard morphometric features. We find that while many features summarizing the overall morphology can be accurately predicted, features related to the topology of arbor shapes, such as maximum branch order, are prone to mistakes (Figure 2d and Figure S16).

While the quantitative analyses described above both suggest that automated reconstruction succeeds in broadly capturing neuronal morphology, including separation of axonal vs. den-dritic branches, they also demonstrate that important differences nevertheless remain between automated and manual traces. Therefore, to robustly analyze anatomical innervation patterns against potential topological mistakes introduced by the automated routine, we develop a 2D arbor density representation (ADR)^12, 35–38^ of axons and dendrites registered to a common laminar axis defined by cortical pia and white matter boundaries. Here, the vertical axis represents the distance from pia and the horizontal axis represents the radial distance from the soma node (Figure 2c). Note that this 2D representation still requires 3D imaging because many branches become undetectable in 2D projected images due to noise (e.g., Figure 1b). Moreover, standardizing the orientations of the brain and the tissue slice is challenging in high-throughput experiments so that the rotation around the laminar axis would be hard to control in 3D representations.

**Figure 2:**
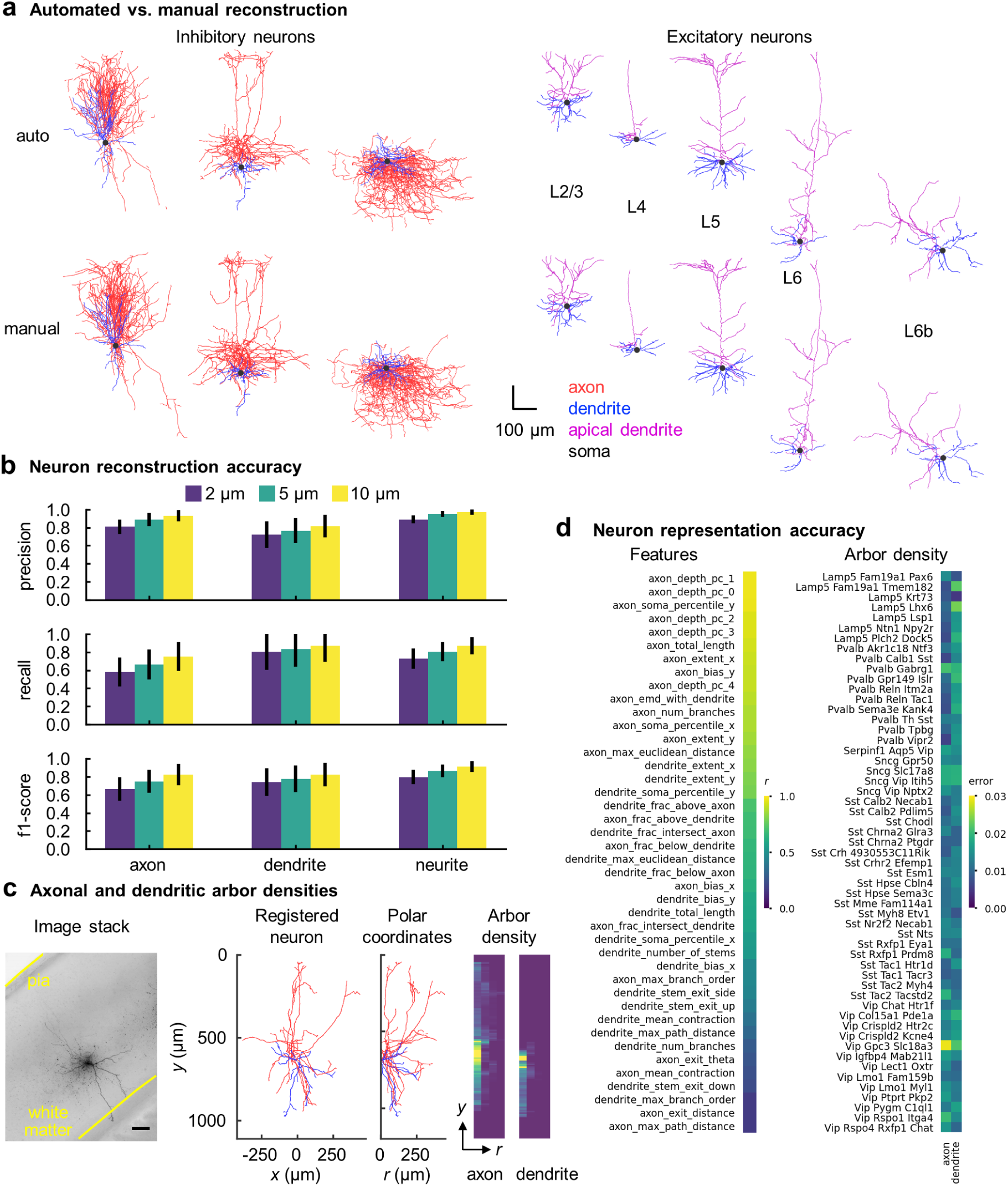
Assessing quality of automated reconstructions. **a**, Automated and manual traces of example test neurons (left, inhibitory neurons; right, excitatory neurons – apical and basal dendrites are assigned for excitatory cells.) **b**, Neuron reconstruction accuracy. Precision, recall, and f1-score values are calculated by comparing automated and manual trace nodes within a given distance (2, 5 and 10 *µm*). Mean values over 340 cells (error bars: standard deviation) are shown for axonal, dendritic and neurite (combined axonal and dendritic) nodes. Scatter plots are shown in Figure S17. **c**, Generation of 2D axonal and dendritic ADRs. Scale bar, 100 *µm*. **d**, *r* values (left) and average root-mean-squared error (right) between automatically vs. manually generated features (left and Figure S16) and ADRs for each t-type (right and Figure S18). ADRs are normalized to have unit norm.

At the level of transcriptomic types, the ADRs calculated from automated reconstructions appear similar to those calculated from manual segmentations based on the root-mean-squared difference between them (Figure 2d). To better quantify this similarity, we compare the performance of the ADR against that of morphometric features^39^ by training classifiers to predict t-types and subclasses (Sst, Pvalb, Vip, Sncg, Lamp5).^5^ We find that the ADR is not statistically significantly worse than the morphometric features in terms of classification accuracy (Boschloo’s exact test, asymptotically exact harmonic mean of p-values over multiple runs:^40^ *p* = 0.81 for t-types, *p* = 0.26 for subclasses, Methods), consistent with Ref.^41^ (Figure 3a,b).

We also test robustness against imperfections due to fully automated tracing by comparing the classification accuracy obtained from automated tracing versus manual tracing on the same set of cells. End-to-end automation appears to perform similarly as manual tracing in cell type prediction based on ADRs (Figure 3c,d) and morphometric features (Figure S19). We finally compare cell type identification based on automatically generated ADRs vs. manually generated morphometric features. We find that they are not significantly different in t-type classification (Boschloo’s test, harmonic mean *p* = 0.72), but the *∼* 5% advantage of manual morphometric features in subclass classification is statistically significant (Boschloo’s test, harmonic mean *p* = 0.01).

Beyond the comparative aspect, these results demonstrate a correspondence between gene expression and the anatomy of local arbors as represented by the proposed registered 2D ADRs, which agrees with previous findings with morphometric features for these cells.^5^ (subclass accuracy of *∼* 79% vs. random at 20%, most abundant label at 47%; t-type accuracy of *∼* 45% vs. random at *∼* 2%, most abundant label at *∼* 8%.) When the transcriptomic type assignments are incorrect, cells are rarely assigned to transcriptomically far-away clusters based on the ADR or morphometric features, as demonstrated by the dominance of the entries around the main diagonal in Figure 3. Note that the rows and columns of these confusion matrices are organized based on the reference taxonomy to reflect transcriptomic proximity (Figures S20 and S21). Therefore, the relative inaccuracy at the t-type level could be attributed to aspects of morphology not captured by the ADR or morphometric features (e.g., synapse locations), or other observation modalities (e.g., physiological, epigenetic) being key separators between closely related t-types.

### Correlates of gene expression and laminar innervation

Having established that registered 2D ADRs are as successful as a standardized, rich set of morphometric features in predicting transcriptomic identity and that ADRs can be generated in a fully automated manner from raw images with only mild loss in performance, we aim to uncover more explicit connections between gene expression and anatomy as captured by the ADR. Since layer-specific axon and dendrite innervations are prominently and reliably captured by the ADR, we study their transcriptomic correlates. We treat the search for genes that are predictive of laminar innervation strength (neurite length innervating a given layer) as a sparse regression problem^42, 43^ (Methods), and focus on 7 t-types whose morphologies are well sampled in our dataset with the help of automated reconstruction (3 Sst, 2 Pvalb, 2 Lamp5 types). That is, we aim to uncover minimal gene sets whose expression can predict the amount of axonal and dendritic innervation of individual laminae as well as the locations of the soma and centroids of the axonal and dendritic trees along the laminar axis. Throughout, we control the false discovery rate (FDR) by applying multiple testing correction (Methods). Tables 1 and S3-S5 summarize these results. We observe that no single anatomical feature is significantly predictable from gene expression for all inhibitory t-types and every studied t-type has at least one significantly predictable anatomical feature. (Only the L4 dendritic innervation strength is significantly correlated with gene expression for cells of type Sst Chodl.) Perhaps more interestingly, we find that the sets of laminae innervation-predicting genes within transcriptomically defined subclasses and t-types are highly reproducible (Table S5) and almost mutually exclusive(Figure 4g). These observations support a connectivity-related organization of cortical cells.^44^ (Multiple discrete and continuous factors of variability may shape neuronal phenotypes^6, 45^ and their dissection may not be possible by studying a subset of non-adjacent t-types.) They also put forth a related question: can gene expression further predict innervation strength of a single layer in a continuum?

**Figure 3:**
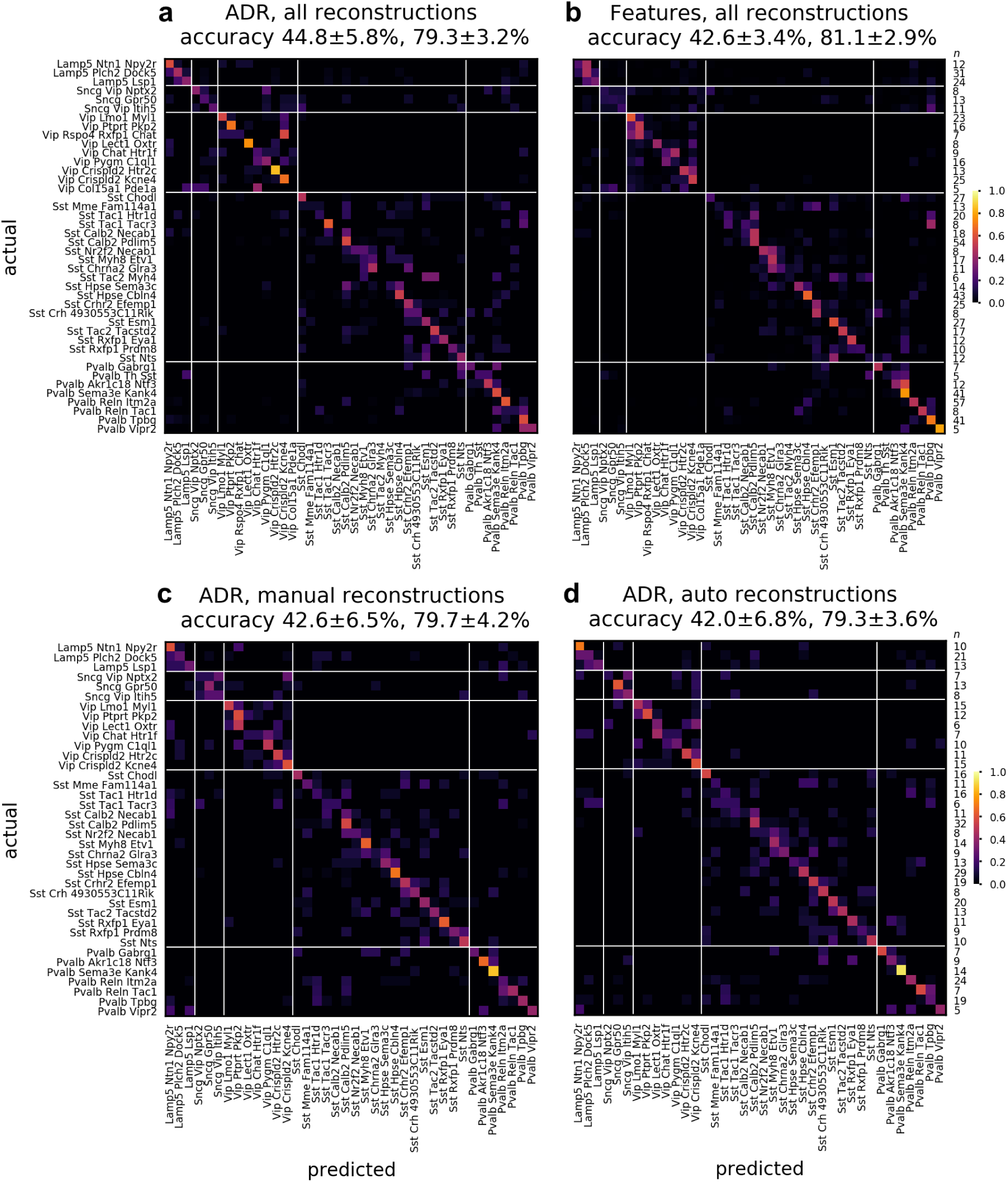
Comparison of cell type classification accuracy based on the ADR vs. a set of classical morphometric features. Confusion matrix for the classification of 42 t-types based on axonal and dendritic ADRs (**a**) and morphometric features (**b**), using a combination of 246 automatically and 501 manually reconstructed cells. Confusion matrix for the classification of 38 t-types based on ADRs, using 488 manually (**c**) and automatically (**d**) reconstructed cells. Accuracy values reported in the headers refer to mean *±* s.d. of the overall t-type and t-subclass classifiers, respectively, across cross-validation folds. Rightmost columns list the number of cells in each t-type (*n*).

**Table 1:** Statistical significance and effect size values for predicting anatomical features from gene expression via sparse linear regression for five different cell types. For each entry, the FDR-corrected *p*-value as calculated by a non-parametric shuffle test is listed. If the value is considered statistically significant at *p ≤* 0.05, the *R*^2^ value is also displayed (*p* / *R*^2^). *p*-values less than or equal to 0.05 and *R*^2^ values larger than or equal to 0.25 are shown in bold. A *p* value of 0 indicates that the calculated *p* value is less than 0.001, the sensitivity of the shuffle test, and less than 0.013 after FDR correction.

|  | Sst Calb2 Pdlm5 | Sst Hpse Cbln4 | Sst Chodl | Pvalb Reln Itm2a |
| --- | --- | --- | --- | --- |
| L1 axon | **0 / 0.40** | 1.000 | 0.637 | **0.023 / 0.13** |
| L2/3 axon | **0.013 / 0.30** | 1.000 | 1.000 | **0 / 0.35** |
| L4 axon | 0.689 | 0.332 | 0.607 | **0 / 0.35** |
| L5 axon | **0.033 / 0.15** | **0.013 / 0.25** | 1.000 | 0.102 |
| L1 dendrite | **0.042 / 0.17** | 1.000 | 0.689 | 0.088 |
| L2/3 dendrite | **0.033 / 0.15** | 1.000 | 0.332 | **0.013 / 0.27** |
| L4 dendrite | 0.697 | **0 / 0.40** | **0.013 / 0.40** | 0.058 |
| L5 dendrite | **0 / 0.19** | 0.102 | 0.393 | **0 / 0.25** |
| soma depth | **0 / 0.26** | **0.023 / 0.21** | 0.246 | **0 / 0.34** |
| axon centroid | **0.023 / 0.16** | 0.210 | 0.058 | **0 / 0.34** |
| dendrite centroid | 0.150 | **0.023 / 0.26** | 0.096 | **0 / 0.39** |
|  | Pvalb Tpbq | Lamp5 Lsp1 | Lamp5 Plch2 Dock5 |  |
| L1 axon | **0.033 / 0.21** | **0.013 / 0.56** | **0.042 / 0.40** |  |
| L2/3 axon | **0.023 / 0.27** | **0.042 / 0.52** | 0.323 |  |
| L4 axon | 0.216 | 1.000 | 0.351 |  |
| L5 axon | 0.058 | 0.135 | 0.081 |  |
| L1 dendrite | **0 / 0.42** | 0.283 | **0 / 0.40** |  |
| L2/3 dendrite | **0.042 / 0.21** | 1.000 | 1.000 |  |
| L4 dendrite | 0.074 | 0.393 | 1.000 |  |
| L5 dendrite | **0.013 / 0.26** | 1.000 | 1.000 |  |
| soma depth | **0.013 / 0.29** | **0.023 / 0.48** | 0.067 |  |
| axon centroid | 0.058 | **0.013 / 0.44** | 0.074 |  |
| dendrite centroid | **0.013 / 0.38** | **0.023 / 0.46** | 0.074 |  |

### Tuning laminar innervation within a cell type: a comparative study

To elucidate this question, we choose a transcriptomically defined subpopulation that is well-sampled in our dataset with the help of automated reconstruction, produces a large effect size in the gene regression study (Table 1), and has been a source of confusion due to its anatomical variability: Sst Calb2 Pdlim5 neurons^23^ represent a transcriptomically homogeneous subset of Martinotti cells, which are inhibitory neurons with L1-innervating axons that gate top-down input to pyramidal neurons.^25, 26^ However, the amount of axon reaching L1 varies widely across cells.^5^ Tables 1 and S5 show that a small set of genes, including genes implicated in synapse formation, cell-cell recognition and adhesion, and neurite outgrowth and arborization,^46–48^ can nevertheless predict the L1-innervating skeletal mass of neurons belonging to this homogeneous population (*R*^2^ = 0.40, *p <* 0.001, non-parametric shuffle test). Since the somata of this population are distributed across L2/3, L4, and L5 (Figure 4b), one potential explanation for this result is that gene expression is correlated with the overall depth location of the cells rather than L1 innervation strength in particular (Figure 4c). Therefore, we repeat the sparse regression study after removing the piecewise linear contribution of soma depth to L1 innervation (linear fit and subtraction for only the L1 innervating subpopulation because the relationship is trivially nonexistent for the non-innervating subpopulation, Methods). We find that the expression levels of a small set of genes are still statistically significantly predictive of L1 innervation: *R*^2^ = 0.31, *p <* 0.001, non-parametric shuffle test. (Repeating with a linear fit and subtraction for the whole population does not change the qualitative result: *R*^2^ = 0.30, *p <* 0.001.)

Next, we obtain a comparative perspective on the L1 innervation result for the Sst Calb2 Pdlim5 subset of Martinotti cells by juxtaposing this result with that for the cells of the Lamp5 Lsp1 type. Somata of these cells are also distributed across multiple cortical layers and their axons have highly variable levels of L1 innervation. Sparse regression again succeeds in finding a small set of genes whose expression level can predict L1 innervation (*R*^2^ = 0.56, *p* = 0.01, non-parametric shuffle test, Table 1). However, it fails to uncover a statistically significant gene set after removing the piecewise linear contribution of soma depth: *R*^2^ = 0.06, *p* = 0.26, non-parametric shuffle test. (Repeating with a linear fit and subtraction for the whole population does not change the qualitative result: *R*^2^ = 0.08, *p* = 0.39.) That is, in contrast to the Sst Calb2 Pdlim5 cells, soma depth almost completely explains the variability in L1 innervation for cells in the Lamp5 Lsp1 population (Figure 4e,f).

**Figure 4:**
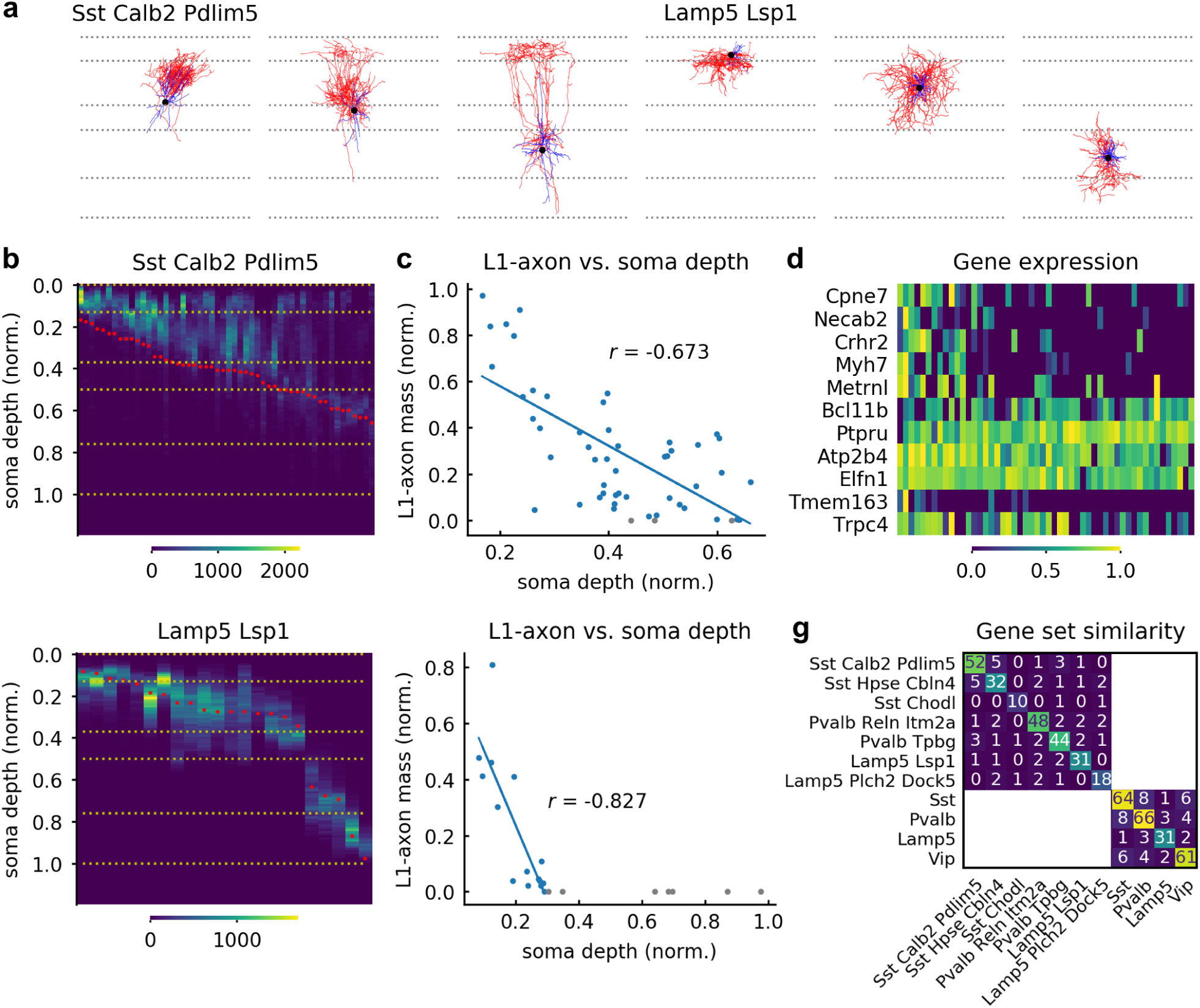
L1 axonal innervation correlates with expression of subset of genes in Martinotti cells. Example neurons of Sst Calb2 Pdlim5 and Lamp5 Lsp1 t-types (**a**). 1D axonal arbor density for the 52 cells in the Sst Calb2 Pdlim5 t-type (**b**) and the 22 cells in the Lamp5 Lsp1 t-type (**e**). (Yellow horizontal dashed lines and red dots indicate cortical layer boundaries and soma depth, respectively). Normalized L1-axon skeletal mass vs. normalized soma depth (0:pia, 1:white matter boundary) for the Sst Calb2 Pdlim5 cells (**c**) and the Lamp5 Lsp1 cells (**f**). Lines fitted to cells with nonzero L1 innervation. Cells whose axons don’t reach L1 are shown in gray. **d**, Gene expression vs. L1-axon skeletal mass for the genes selected by the sparse regression analysis. (L1-axon mass decreases from left to right.) **g**, Similarity matrix for the sets of laminae-predicting genes within transcriptomic types and subclasses. (See Table S5.) Each entry denotes the number of genes in the intersection between the corresponding row and column.

Lastly, we consider the possibility that the cells whose axons do not reach L1 are simply irrelevant for this study and bias the statistics. (Axons of 3 out of 52 cells in the Sst Calb2 Pdlim5 population, and 7 out of 22 cells in the Lamp5 Lsp1 population do not reach L1.) We repeat the above comparison after removing the cells whose axons don’t reach L1 altogether from this study. Sparse regression still uncovers a statistically significant relationship between L1 innervation strength and a set of genes for the Sst Calb2 Pdlim5 population after removing the linear contribution of soma depth (*R*^2^ = 0.28, *p <* 0.001). In contrast, it again fails to find a statistically significant relationship for the Lamp5 Lsp1 population (*R*^2^ = 0.05, *p* = 0.39).

To summarize, while axons of Lamp5 Lsp1 cells appear to shift along the laminar axis according to their soma location within the cortical depth, soma location does not seem to dictate the axonal L1 innervation of Sst Calb2 Pdlim5 neurons, whose strength can nevertheless be predicted by gene expression. For both of these t-types, the automated reconstruction pipeline increased the sample size by more than 60% (Sst Calb2 Pdlim5: 63%, Lamp5 Lsp1: 69%), empowering the statistical analysis pursued here. Similarly, the sample counts for the t-types studied in Table 1 increased between 48% and 138%. (The increase over the whole dataset is 50%, from 543 to 813 cells.) Since t-types correspond to leaf nodes of the cell type hierarchy, their sample sizes are much smaller than the subclass-level counts. Therefore, automated reconstruction can be beneficial both by capturing more of the biological variability in single cell morphologies of populations at the finest level of transcriptomically defined taxonomies and by enabling cross-validation schemes similar to the ones pursued here.

## Discussion

While classification of neuronal cell types is increasingly based on single cell and nucleus genomic technologies, characterization of neuron morphology – a classical approach – captures an aspect of neuronal identity that is stable over long time scales, is intimately related to connectivity and function, and can now be connected with genomic attributes through the use of simultaneous profiling techniques such as Patch-seq. Nevertheless, light microscopy based methods of neuronal reconstruction often inadequately reproduce the determinant attributes of morphological signature, especially in high-throughput settings. Here, we have presented an end-to-end automated neuronal morphology reconstruction pipeline for brightfield microscopy, whose simple setup supports flexible, single or multimodal, characterization protocols. We have also proposed an arbor density representation as a descriptor of cortical neuronal anatomy that is robust against noise in high-throughput imaging scenarios as well as mistakes of automated reconstruction. Its success suggests that detailed morphological reconstructions may ultimately not be necessary if the only aim is inferring the cell type label.

Through the use of sparsity arguments and statistical testing, we demonstrated that this pipeline can help reveal relationships between gene expression and neuronal anatomy, where a large number of anatomical reconstructions enables accurate inference in the presence of a large gene set. As an application, we studied the correlation between gene expression and laminar innervation on a Patch-seq dataset of cortical neurons^5^ and showed that the gene correlates of different innervation patterns have little overlap across transcriptomically defined subpopulations. While the same program can potentially also address the relationship between morphological and electrophysiological properties of neurons, the accuracy of automated reconstructions should further improve for use in detailed compartmental models.^49^

Finally, we focused on axonal innervation of L1 by a transcriptomically defined subpopulation of Somatostatin-expressing Martinotti cells. We found that the innervation strength is relatively weakly correlated with soma depth for this cell type, but not all types. Moreover, a subset of genes can predict the remaining variability in the innervation strength after the effect of soma depth is removed, suggesting a control mechanism beyond simple shifting of the morphology within the cortical depth for this cell type. Considering that neurons in this population are thought to gate top-down input to cortical pyramidal neurons,^25, 26^ this result suggests tuning of innervation strength in a continuum within the discrete laminar organization of the mouse cortex,^45, 50–52^ potentially to improve task performance of the underlying neuronal network.

From a segmentation perspective, we believe our work represents a significant step forward as the first study to produce hundreds of automatically reconstructed morphologies obtained from the brightfield microscope (Fig. S1-S11). As demonstrated in the main text, these cortical neuron morphologies are statistically indistinguishable from their manually generated counterparts in certain aspects (e.g., cell type identification), but not in many others (e.g., arbor topology). Indeed, much further improvement is needed to achieve complete and accurate tracing of neurons. Nevertheless, advances in computer vision algorithms and computing infrastructure that can support complicated models and large datasets suggest that qualitative improvements may be within reach in the next few years. Larger training sets will improve generalization, enable the use of larger image contexts and the effective tuning of more parameters (e.g., the use of the popular transformer architecture.^53, 54^) The presented method can increase the speed of manually verified trace generation. In addition, existing manually traced neurons without transcriptomic characterization (e.g., Ref.^39^) can still be useful in training the segmentation model. Finally, while voxel-based loss functions, such as the one used in our model, are easier and faster to train, single voxel mistakes can change the connectivity due to the filamentous appearance of the arbor under the light microscope. Therefore, topology-aware objective functions^55, 56^ can improve the topological accuracy of the segmentations, a relative weakness of the proposed model. If perfect segmentation is required, we expect a human expert to remain in the loop in the near future, primarily to verify the accuracy of the branching points.

## Methods

### Dataset

The dataset profiling local morphology and transcriptome of GABAergic mouse visual cortical neurons was generated as part of the Patch-seq recordings described in Ref.^5^ This dataset includes 2,341 cells with transcriptomic profiling and high resolution image stacks, where the brain sections were imaged on an upright AxioImager Z2 microscope (Zeiss) equipped with an Axiocam506 monochrome camera. Tiled image stacks of individual cells were acquired with a 63*×* objective lens (Zeiss Plan-Apochromat 63*×*/1.4 NA or Zeiss LD LCI Plan-Apochromat 63*×*/1.2 NA) at an interval of 0.28 *µm* or 0.44 *µm* along the Z axis. Individual cells were manually placed in the appropriate cortical region and layer within the Allen Mouse Common Coordinate Framework (CCF)^57, 58^ by matching the 20X image of the slice with a “virtual” slice at an appropriate location and orientation within the CCF. 1,259 cells were removed from the dataset either because they were mapped to nearby regions (instead of visual cortex) or because their images had incomplete axons. 543 of the remaining 1,082 bright-field image stacks of biocytin-filled neurons were reconstructed both manually and automatically, and this set is used for training and testing of the t-type classification algorithms and for error/*R*^2^-value quantification. The remaining 539 cells were reconstructed only automatically. To ensure the quality of scientific results presented in Table 1 and Figure 4, we excluded 118 images with multiple neurons in the field of view and 151 images that failed at different stages of the pipeline (missing pia/white matter annotations, annotation-related failed upright transformation, reconstruction failing visual inspection) from those analyses. Finally, 16 cells in the manually and automatically reconstructed population and 20 cells in the automatically-only reconstructed population were not used for analyses involving t-types because these cells were deemed to not have “highly consistent” t-type mappings in Ref.^5^

### Volumetric training data generation from skeletonized morphologies

Segmentation of neuronal morphologies from 3D image stack requires voxel-wise labels while manual reconstructions specified by traces only provide a set of vertices and edges corresponding to the layout of the underlying morphology. We developed a topology-preserving fast marching algorithm to generate volumetric ground truth using raw image stacks and manual traces by adapting a fast-marching based segmentation algorithm^13, 59^ initialized with trace vertices to segment image voxels. This segmentation should be consistent with the layout of morphology traces, without introducing topological errors (e.g., splits, merges, holes). We ensured this by incorporating simple point methods in digital topology^60^ into the segmentation algorithm. (i.e., the proposal generated by fast marching is disallowed if the proposed voxel value flip changes the underlying topology.) We noticed that the soma can be incompletely filled by the fast marching algorithm. Therefore, we treated the soma region separately and used a sequence of erosion and dilation operations followed by manual thresholding to achieve complete labeling. Each voxel was labeled as axon, dendrite, soma, background.

### Neural network architecture and training

We used a 3D U-Net convolutional neural network^28^ to perform multi-class segmentation (i.e. each voxel is assigned a probability of belonging to classes specified in the label set). We trained two separate models using raw images and volumetric ground truth; one with 51 inhibitory neurons, and another with 75 excitatory neurons from mouse cortex. The weights of the excitatory model were initialized with those of the trained inhibitory model, except for the classification layer. The U-Net architecture consists of a contracting path to capture context and a symmetric expanding path that enables precise localization and has been shown to achieve state-of-the-art performance in many segmentation tasks.^27, 28^ Building on previous work,^16^ we developed an efficient Pytorch implementation that runs on graphical processing units (GPUs). To address GPU memory constraints, during each training epoch the training set was divided randomly into subsets of 3 stacks. Training proceeded sequentially using all subsets in an epoch, and data loading time did not exceed 10% of the total training time. We trained the model on batches of 3D patches (128 *×* 128 *×* 32 px^3^, XYZ), which were randomly sampled from the subset of 3 stacks loaded into memory. All the models were trained using the Adam optimizer^61^ with a learning rate of 0.1. Training with a GeForce GTX 1080 GPU for *∼* 50 epochs took *∼* 3 weeks. Since the neuron occupies a small fraction of the image volume, we chose patches that contained at least one voxel belonging to the neuron. To add salient, negative examples to the training set, we also included a number of patches with bright backgrounds produced by staining artifacts and pial surface. To enable the model to generalize from relatively small number of training examples and improve the segmentation accuracy, we augmented training data by 90*^◦^* rotations and vertical/horizontal flips in the image (XY) plane.

### End-to-end neuron reconstruction pipeline

An end-to-end automated pipeline combined segmentation of raw image stacks into soma, axon, and dendrite structures with post-processing routines to produce a swc file. Segmentation with trained models using a single high-end GPU takes *∼*5.8 min per Gvoxel, or *∼*186 min for average 32 Gvoxel image stack. Our pipeline had access to 16 GPU (NVIDIA Titan X) cards. Even though the model trained using inhibitory neurons generalized well on all types of neurons, we found that the model trained using excitatory neurons improved the axon/dendrite assignment accuracy on excitatory neurons. Segmentation was post-processed by thresholding the background, followed by connected component analysis to remove short segments, skeletonization, and converting to a digital reconstruction in the swc format.

### Axon/dendrite relabeling

Since the initial segmentation by the UNet-based neural network assigns every foreground voxel one of {soma, axon, dendrite} based on local context defined by the patch size, it is prone to occasional errors, particularly in distinguishing between axon and dendrite. These errors propagate to the labeling of nodes in the tree representation. In order to improve this initial node labeling, we developed an error correction approach which utilizes the initial segmentation to make a decision based on a larger spatial context.

We trained a secondary neural network model to predict the labels based on features calculated using the raw image stack and the initial trace. First, for each connected component in the skeleton of the initial segmentation, we identified the longest path from the node closest to soma and calculated features on the node set defining that path: (i) The first 6 features are 1D arrays of image brightness values calculated for every node in the set at different spatial scales using spherical kernels of varying radii (1, 2, 4, 8, 16, 32). (ii) A second set of 6 features are 1D arrays of neurite tortuosity values calculated for every node in the set as ratios of path and Euclidean distances between each node and its n-th neighbor away from the soma (n = 4, 8, 16, 32, 64, 128). (iii) Two additional features are 1D arrays of node type of every node in the set and a single number representing distance of the closest node to the soma. For each 1D array, we used the first 2048 nodes and zero-padded shorter arrays to have a uniform array size.

The neural network architecture consists of three arms, two arms have two convolutional layers with 4 and 8 7x3 filters followed by 4x1 max pooling, a fully connected layer and a dropout layer each. These arms process feature sets (i) and (ii) above by stacking the sets of 6 1D arrays along a second dimension. The third arm processes feature set (iii) and has two convolutional layers with 4 and 8 7x1 filters followed by 4x1 max pooling, a fully connected layer and a dropout layer. The outputs of these three arms are concatenated with the ‘distance to soma feature’. Finally, the concatenated hidden feature map is processed by two fully connected layers to produce a single scalar softmax output indicating the inferred label type. The network model was trained using examples from the training dataset of the semantic segmentation model, and is used to relabel the neuron traces during postprocessing. Averaging predicted label type over the 9 longest paths improved relabeling accuracy for connected components longer than 2048 nodes.

### Connecting disconnected segments

Due to a combination of staining artifacts and limitations of brightfield imaging, the initial reconstruction is often characterized by multiple disconnected subtrees. We introduced artificial breaks to manual traces to train a random forest classifier to predict whether nearby pairs of connected components should be merged. First, we find all pairs of segment (subtree) end points that are located within a certain distance. Next, for each pair of end points, we calculate the Euclidean distance and four collinearity values which measure the segments’ orientation relative to each other. Specifically, for each end point, we calculate two vectors representing segment end orientation at two different spatial scales (i.e., the orientation of the branch terminating at that end point). The collinearity values are the dot products of each of these vectors with the vector between end points of the pair. Finally, for each end point in the pair we also calculate the above features for the closest four other end points. As a result, for every pair of end nodes we have a total of 45 features. Only segments of the same type, axon or dendrite, are considered for merging.

### Additional postprocessing

We passed the reconstructions through a series of additional post-processing for extraneous cell/artefact removal, down sampling, node sorting and pruning. We used quality control by a human to check for the presence of disconnected branches of extraneous cells that were not removed during postprocessing. We excluded samples that did not pass this quality control if they did not have a manual trace. For samples with a manual trace, we used a neighborhood of the manual trace to simulate manual removal of extraneous cells by masking with that neighborhood. We excluded these samples from reconstruction accuracy quantification and used them only for cell type classification. For excitatory neurons (Fig. 2a, and Ref.^34^), we trained a random forest classifier to identify apical dendrite segments in excitatory auto-trace reconstructions. The classifier was trained on geometric features that distinguish apical dendrite segments from the basal dendrite (e.g. upright distance from soma). This classifier achieved a mean accuracy of 85% percent across 10-fold cross validation.

### Morphometric feature calculation

Reconstructions were transformed to an upright position that is perpendicular with respect to pia and white matter surfaces. Morphometric features as described in^39^ were calculated using the skeleton_keys python package. Following Ref.,^39^ z-dimension features were not included.

### Arbor density generation of axons and dendrites

We represented axonal and dendritic morphologies as 2D density maps registered to a common local coordinate axis using the pia/white matter boundaries. First, we applied upright transform to the reconstructed neuron followed by the correction of z-shrinkage and the variable tilt angle of the acute slice.^39^ Next, adapting our previous work^12, 38^ we conformally mapped pia and white matter boundaries to flat surfaces, calculated a nonlinear transformation on the whole tissue by a least-square fit to pia/white matter mappings, and applied this transformation to the morphology trace. Finally, we used the registered trace to generate a 2D density representation. The polar axes representing the cortical depth and the lateral distance from the soma (Figure 2c) make this representation invariant to rotations around the laminar axis. We calculated these maps separately for axons and dendrites. We downsampled these maps to 120px *×* 4px images with a pixel size of 8*µm ×* 125*µm* to be robust to minor changes in morphology. We normalized the intensity by the lateral area corresponding to each pixel so that each pixel value represents local arbor density.

### Assessing neuron reconstruction accuracy

To assess the quality of automated neuron reconstructions, we used manual reconstructions as the ground truth. We quantified the correspondence of trace nodes, as described in the main text, to evaluate the accuracy of the trace layout. We calculated precision, recall and f1-score metrics at three distances (2, 5, and 10 *µm*), and reported mean and standard deviation values for the test set of 340 neurons (all samples that have manual traces excluding the ones used for training models or required masking). In addition, we evaluated the accuracy of neuron morphology representations, morphometric features and ADRs, for the same set. After realizing that the image coordinates used for manual vs. automated tracing were nonlinearly warped with respect to each other for one cell (penultimate cell in Figure S10), we removed that cell from the single cell-level node correspondence, ADR, and feature-based comparison studies. (It was used in all cell typing studies.) We reported the Pearson correlation coefficient *r* for each morphometric feature. We calculated the average root-mean-squared error per t-type between normalized axon/dendrite ADRs derived from automated and manual reconstructions.

### Supervised classification

Supervised classification using morphometric features was performed by training a random forest classifier implemented in the scikit-learn Python package^62^ using 5-fold cross-validation. This was repeated 20 times. Supervised classification using ADRs was done by training a feed-forward neural network classifier using our Pytorch implementation. The network architecture consists of a convolutional layer with 7x3x1 filters, a 4x1x1 max pooling layer, a convolutional layer with 7x3x1 filters, a 3x1x1 max pooling layer, a layer that concatenates the hidden features with the soma depth value, and a fully connected layer with the number of units corresponding to the number of classes. Each convolutional layer uses the rectified linear function as the non-linear transformation.

Since the depth locations of neurons vary within the cortex, we introduced a novel type of data augmentation based on simulation of cell type-dependent neuronal shift along the laminar axis. Namely, for each t-type we calculated the range of soma depth variations, and applied a random shift within that range to the input soma depth value, as well as the corresponding shift to the ADR intensity in the laminar direction. This cell type-dependent random shift of the ADR and the soma depth together with a modulation of the intensity values of the ADR improved the accuracy of classification.

The networks were trained using the cross-entropy loss function and the Adam optimizer with a learning rate of 0.001. Training using 10-fold cross-validation with GeForce GTX 1080 GPU for 50,000 epochs took *∼* 24 h. A set of 246 automatically and 501 manually reconstructed cells was used for training classifiers shown in Figure 3a and b, and a set of the same 488 automatically and manually reconstructed cells were used for Figure 3c,d and Figure S19. Both sets included only cells from t-types with at least 5 cells. Confusion matrices, mean and standard deviation of accuracy across cross-validation folds were reported.

We performed Boschloo’s exact test on 2x2 contingency tables where columns/rows store total numbers of correct and incorrect predictions for two given classifiers. When comparing ADR-based to morphometric feature-based classifiers, we calculated the contingency table for each of the 20 repetitions used in the feature-based classifier study. We calculated the p-value of one-sided Boschloo’s test to evaluate the null hypothesis of ADR-based accuracy being less than feature-based accuracy. To aggregate the 20 p-values, we used Ref.^40^ and the Python implementation at https://github.com/benjaminpatrickevans/harmonicmeanp to report the p-value of the asymptotically exact harmonic mean p-value test for t-type and subclass predictions.

#### Cophenetic agreement

To take the hierarchical organization of transcriptomically defined mouse cortical cell types^23^ into account when evaluating the accuracies of the different classification tasks, we defined resolution index per cell as the scaled height of the closest common ancestor of assigned and predicted labels in the t-type hierarchical tree^63^ (Figures S20 and S21). Accordingly, the resolution index for a correctly classified t-type (“leaf node” label) is 1. In the worst case, the closest ancestor for an assigned and predicted label can be the root node of the taxonomy, which corresponds to a resolution index of 0. We report the mean and s.e.m. values for this measure of cophenetic agreement between true and predicted assignments for each cell type in Figures S20 and S21.

### Sparse feature selection analysis

Following Ref.,^64^ a set of 1,252 genes were used for this analysis. This set was obtained by excluding genes if they satisfy any of the following criteria: (1) they are highly expressed in non-neuronal cells, (2) they have previously reported sex or mitochondrial associations, and (3) they are much more highly expressed in Patch-seq data vs. Fluorescence Activated Cell Sorting (FACS) data (or vice versa) and therefore may be associated with the experimental platform^5^. Further, we removed gene models and some other families of unannotated genes that may be difficult to interpret. We also used the *β* score, a published measure to evaluate the degree to which gene expression is exclusive to t-types,^65^ to exclude genes expressed broadly across t-types. Gene expression values were CPM normalized, and then log*_e_*(*•* + 1) transformed for all the downstream analyses.

A set of 777 neurons was used for the feature selection analysis where automated reconstructions comprise *∼* 32% of this set (*∼* 44% of the subset of 7 t-types studied in Figure 4). Every neuron in the dataset was characterized by the expression levels of the set of 1252 genes, and their axonal and dendritic 1D arbor density representations were organized into two 120 *×* 1 vectors. For each neuron, axon/dendrite layer-specific skeletal masses normalized by total skeletal mass were calculated to quantify layer-specific innervation for axonal and dendritic morphologies, and axon/dendrite centroids were calculated to characterize laminar position of the morphology. To select a small subset of genes that are responsible for the variability in individual anatomical features within each transcriptomic type or subclass, we solved the Lasso regression problem^66^ (LassoCV command in the scikit-learn library^62^) for each anatomical feature using the cells in that transcriptomic set. We analyzed only the sets corresponding to the types and subclasses with at least 20 cells. Briefly, let *K_t_* denote the number of cells of type or subclass *t* for which we have both anatomical features *y_t_* (a *K_t_ ×* 1 vector) and gene expression values *X_t_* (a *K_t_ × N* matrix). We solve

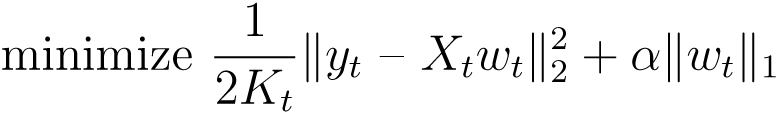

by performing nested 5-fold cross-validation. For each cross-validation fold, we passed the training set into LassoCV which performed another splitting of the data to determine the hyperparameter *α* and the set of selected genes. We selected the 10 genes with maximum absolute weight values and calculated the coefficient of determination, *R*^2^, for the test set. Finally, we selected the 10 most frequent genes across the 5 folds and calculated the mean test *R*^2^ value. To evaluate statistical significance, we shuffled the rows of the gene expression matrix *X_t_* 1000 times and used the same procedure to calculate mean test *R*^2^ value for each shuffled run. We calculated the one-sided *p*-value as the fraction of shuffled runs with *R*^2^ values greater than or equal to the true *R*^2^. Finally, we performed multiple testing correction of p-values using Benjamini-Yekutieli method^67^ to control the false discovery rate (multipletests command in the statsmodels library^68^). We report resulting p-values and test *R*^2^ values for t-types in Table 1 and for t-types and subclasses in Tables S3 and S4.

## Data availability

Transcriptomic and morphological data supporting the findings of this study is available online at https://portal.brain-map.org/explore/classes/multimodal-characterization (“Neurons in Mouse Primary Visual Cortex”). Additional dataset of automated morphological reconstructions is available at https://github.com/ogliko/patchseq-autorecon.

## Code availability

Code pertaining to this study as well as the trained neural network model for automated segmentation are available at https://github.com/ogliko/patchseq-autorecon and https://github.com/rhngla/topo-preserve-fastmarching. Morphometric features are calculated using the skeleton_keys package at https://skeleton-keys.readthedocs.io.

## Acknowledgements

We wish to thank the Allen Institute for Brain Science founder, Paul G Allen, for his vision, encouragement and support. We thank Michael Hawrylycz and Claire Gamlin for helpful feedback on the manuscript. This work was supported by the National Institute Of Mental Health of the National Institutes of Health under award numbers 1U01MH114824-01 and 1RF1MH128778-01. The content is solely the responsibility of the authors and does not necessarily represent the official views of the National Institutes of Health.

## Supplementary Material

**Figure S1:**
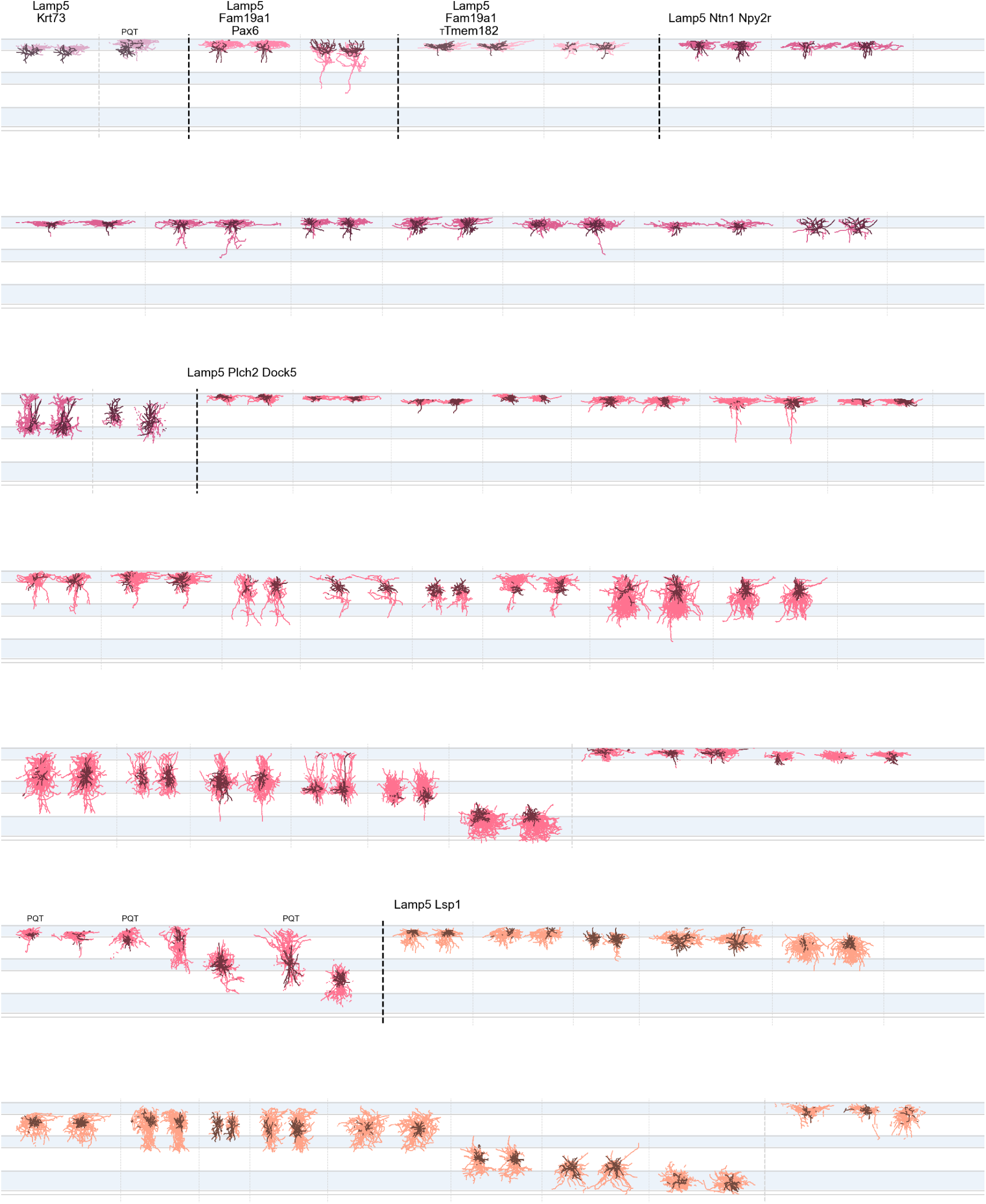
Morphological reconstructions of inhibitory neurons ordered by t-type – 1 of 11. 543 neurons have both automated and manual reconstructions, 270 - only automated ones. For each t-type, cells with both automated and manual reconstructions are shown first, separated by faint dashed lines, followed by cells that are reconstructed only automatically. Dendrites and axon are in darker and lighter colors, respectively. Best viewed digitally. PQT: poor quality transcriptomic characterization – not used for t-type related analyses. T: cells used in segmentation model training.

**Figure S2:**
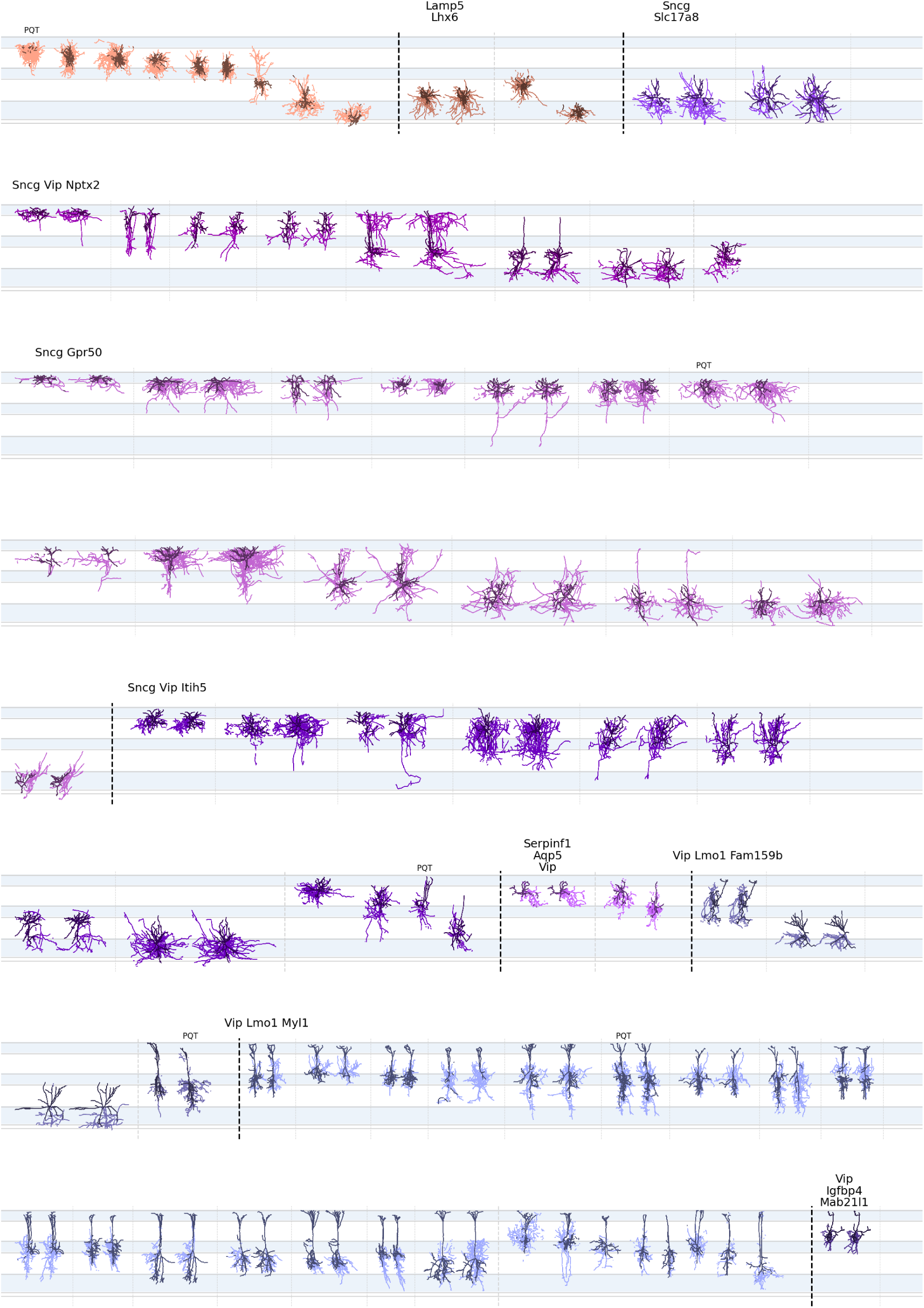
Morphological reconstructions of inhibitory neurons ordered by t-type – 2 of 11. Please refer to the caption of Figure S1.

**Figure S3:**
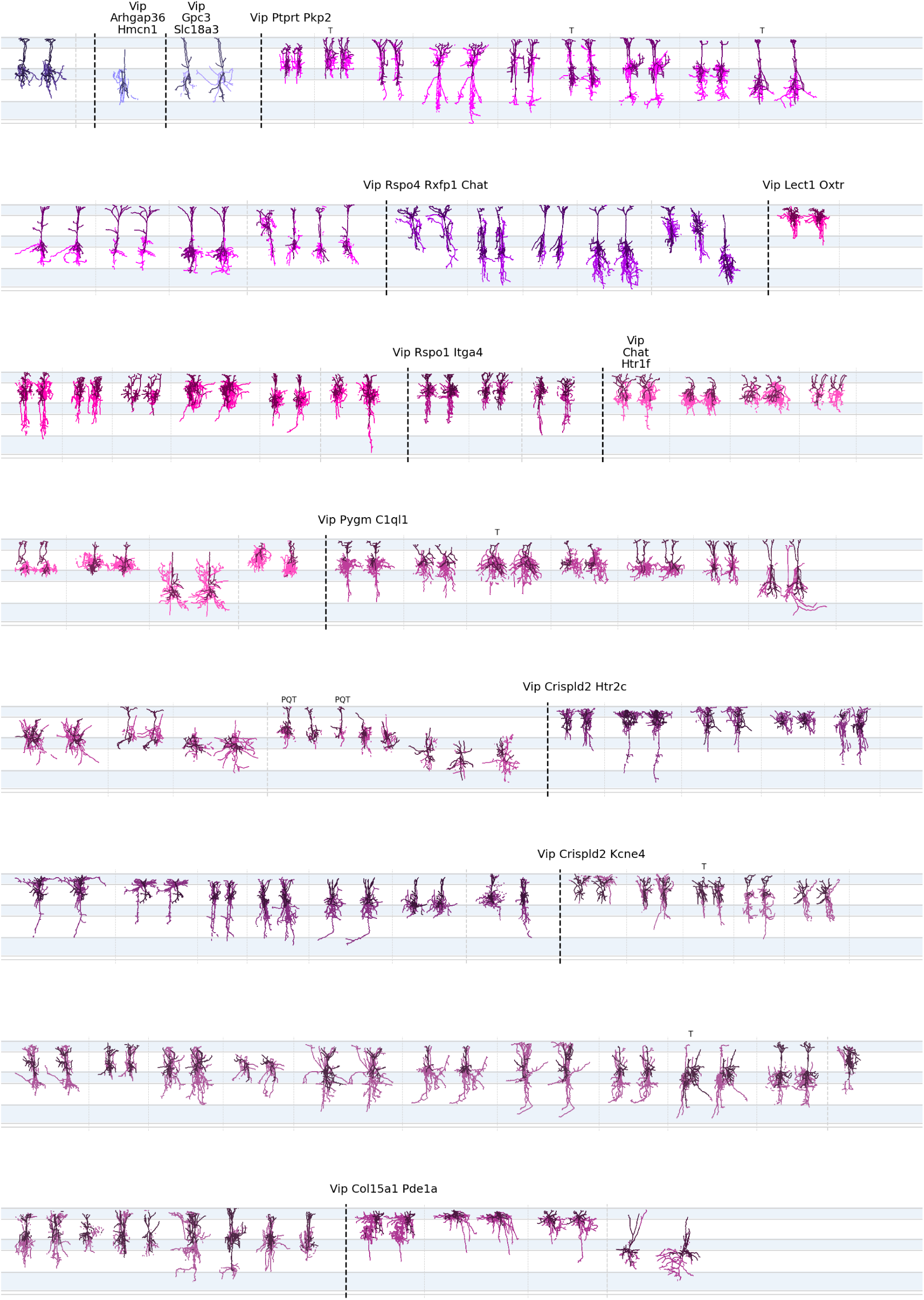
Morphological reconstructions of inhibitory neurons ordered by t-type – 3 of 11. Please refer to the caption of Figure S1.

**Figure S4:**
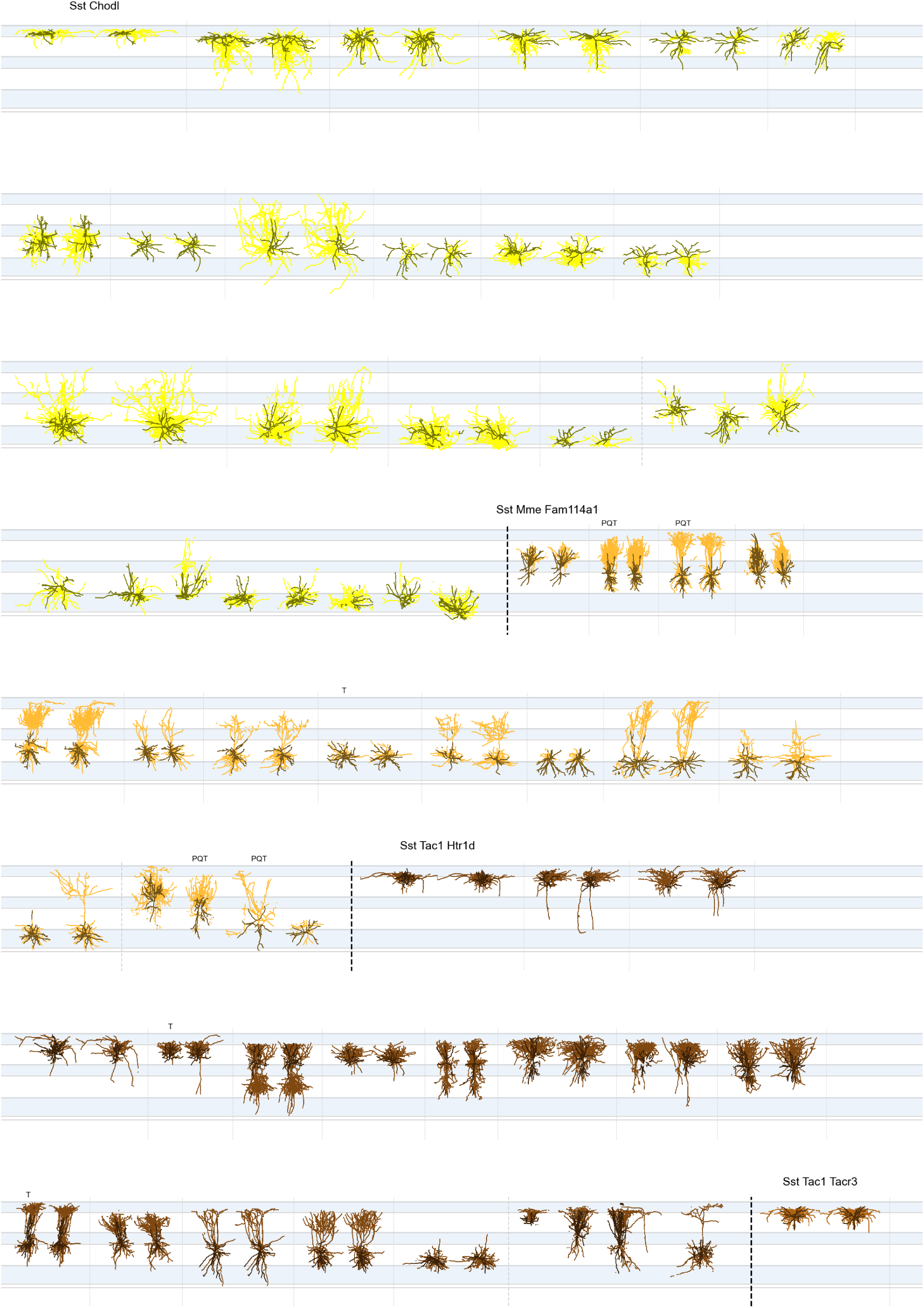
Morphological reconstructions of inhibitory neurons ordered by t-type – 4 of 11. Please refer to the caption of Figure S1.

**Figure S5:**
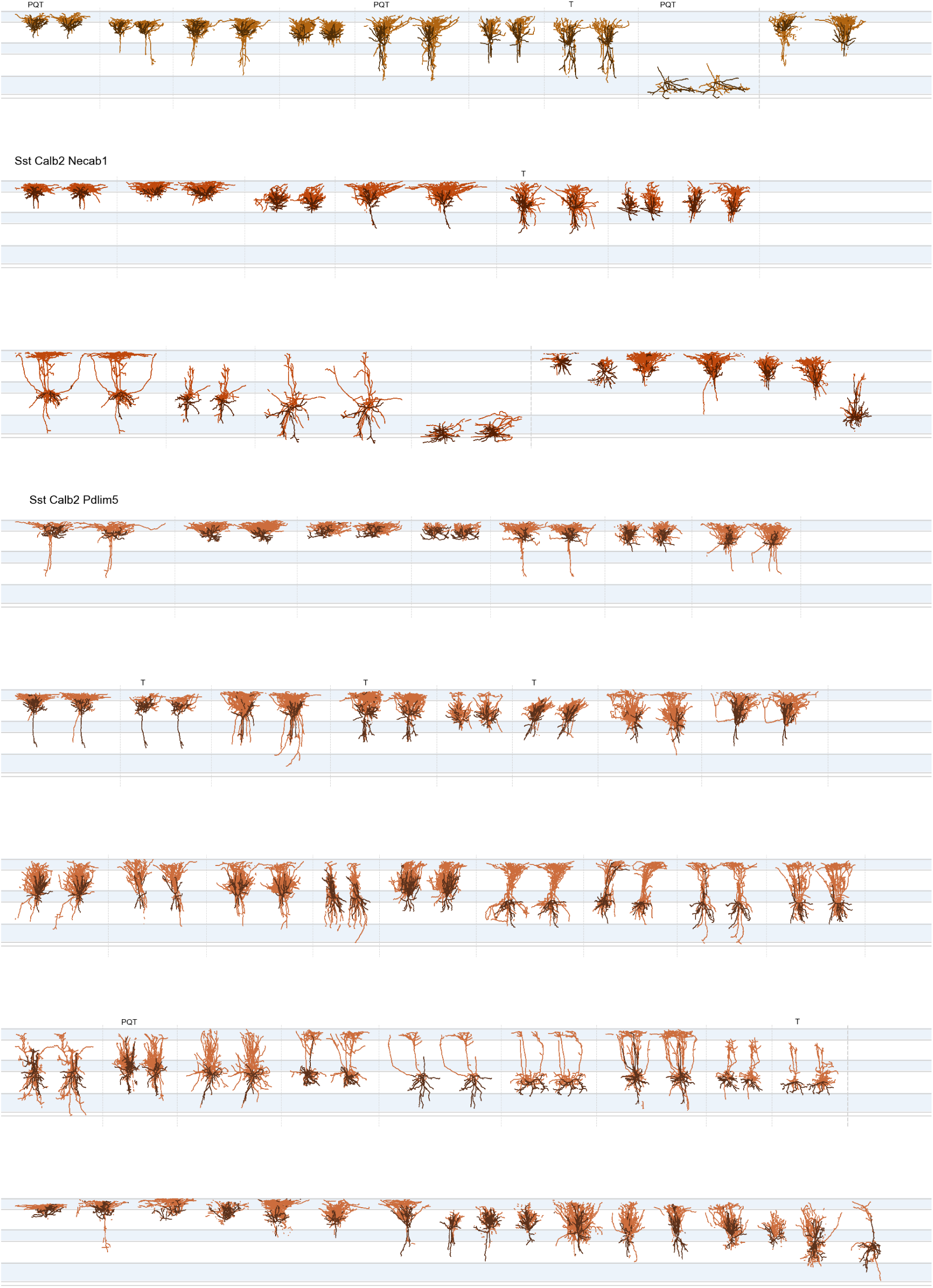
Morphological reconstructions of inhibitory neurons ordered by t-type – 5 of 11. Please refer to the caption of Figure S1.

**Figure S6:**
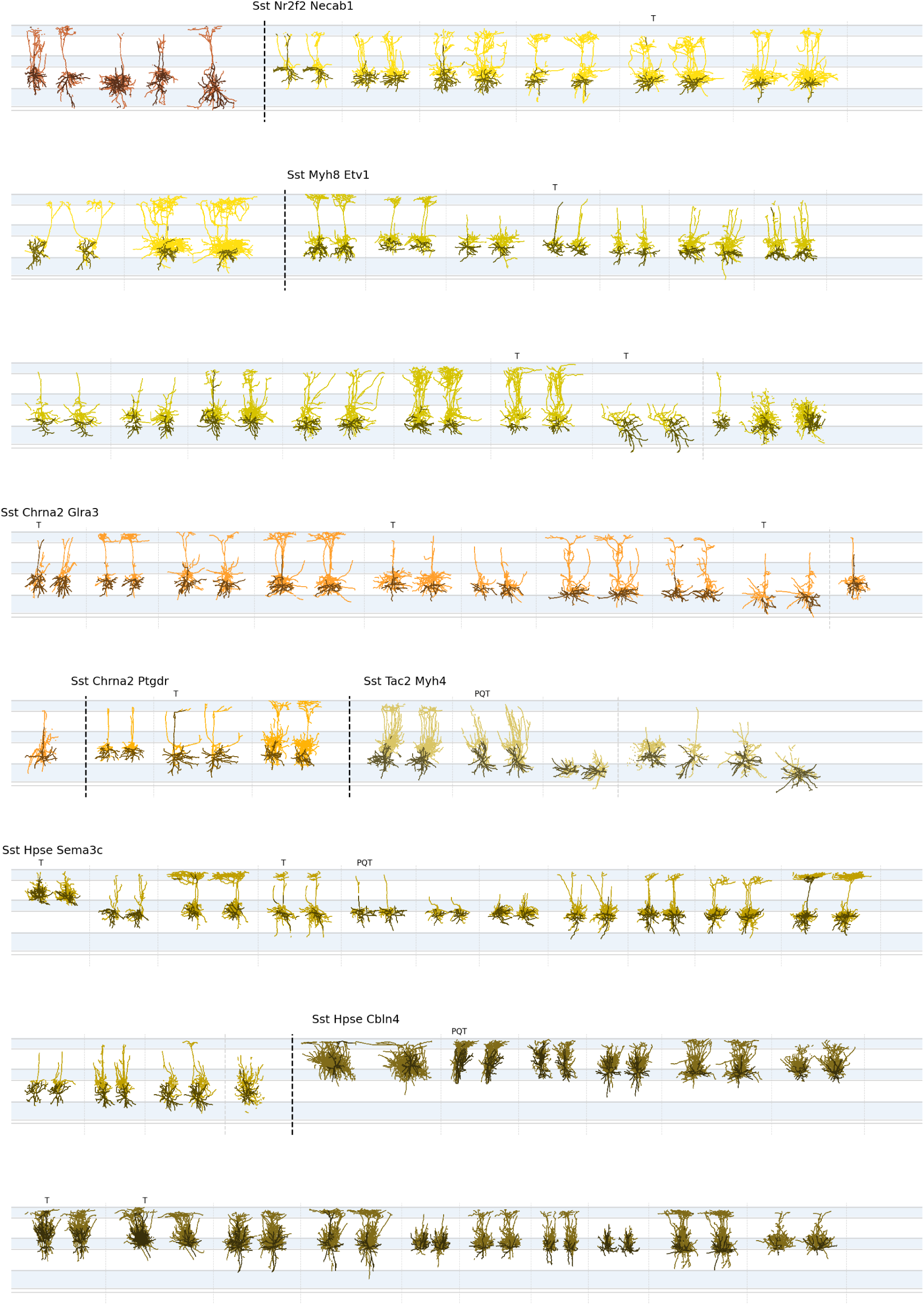
Morphological reconstructions of inhibitory neurons ordered by t-type – 6 of 11. Please refer to the caption of Figure S1.

**Figure S7:**
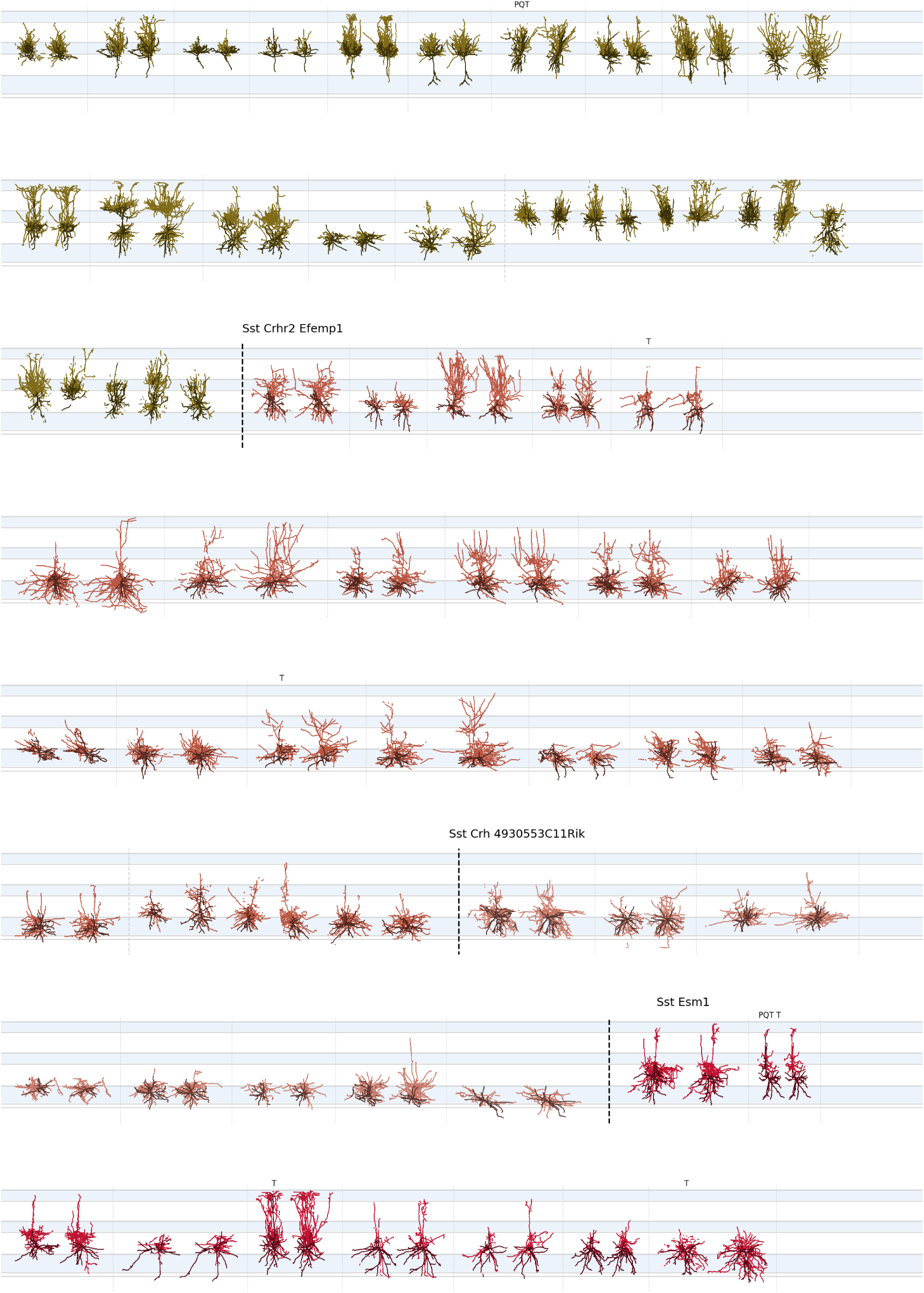
Morphological reconstructions of inhibitory neurons ordered by t-type – 7 of 11. Please refer to the caption of Figure S1.

**Figure S8:**
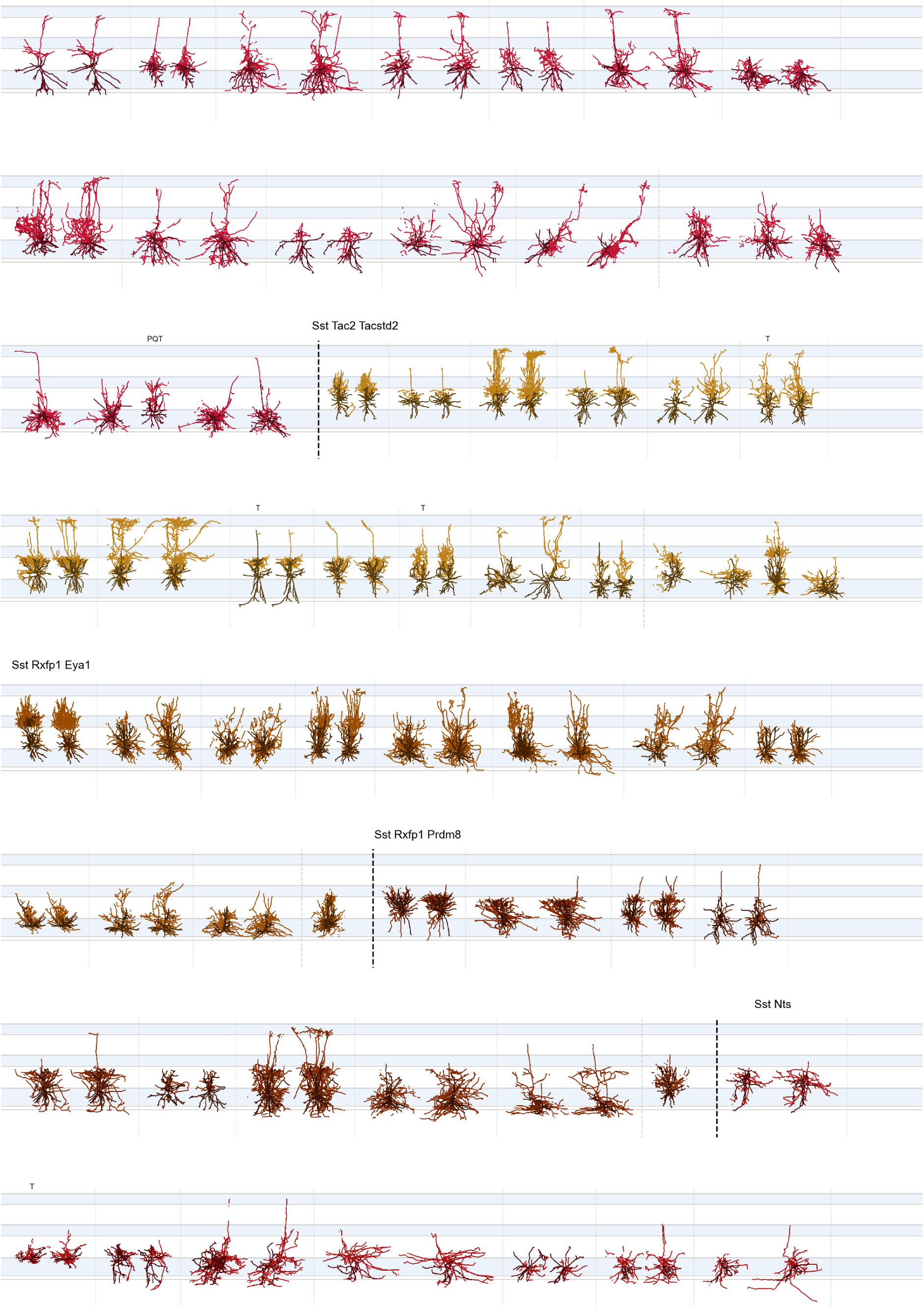
Morphological reconstructions of inhibitory neurons ordered by t-type – 8 of 11. Please refer to the caption of Figure S1.

**Figure S9:**
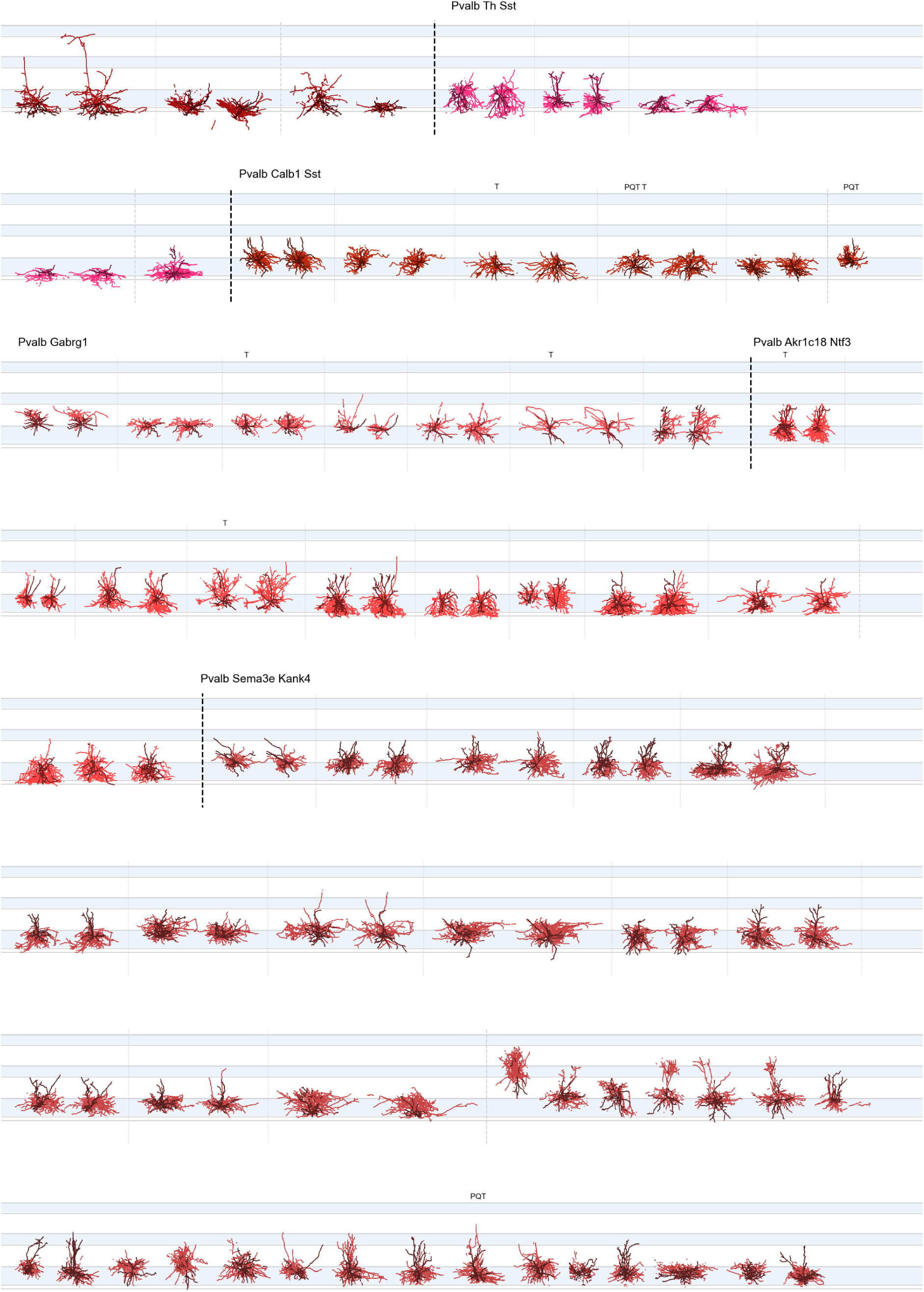
Morphological reconstructions of inhibitory neurons ordered by t-type – 9 of 11. Please refer to the caption of Figure S1.

**Figure S10:**
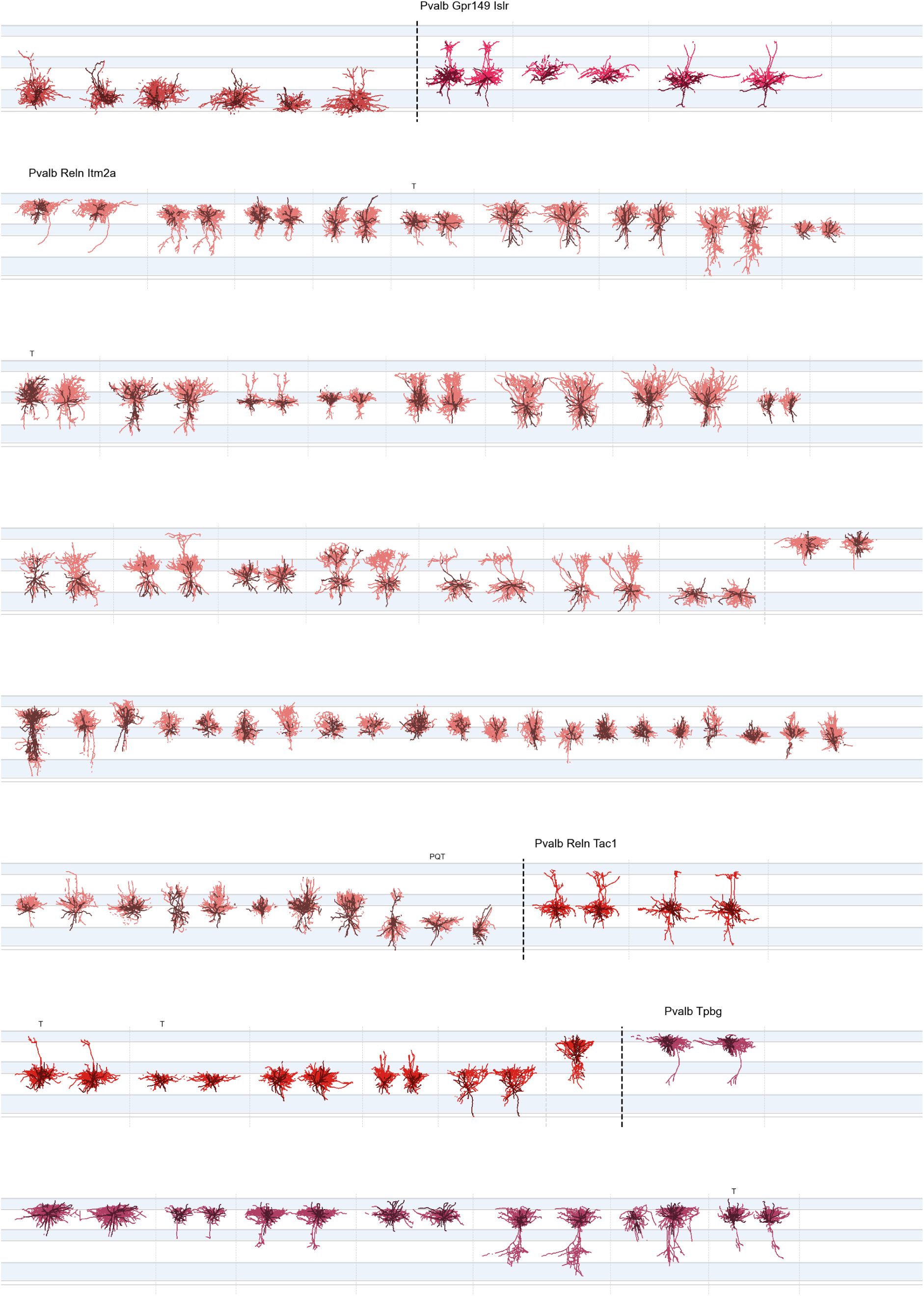
Morphological reconstructions of inhibitory neurons ordered by t-type – 10 of 11. Please refer to the caption of Figure S1.

**Figure S11:**
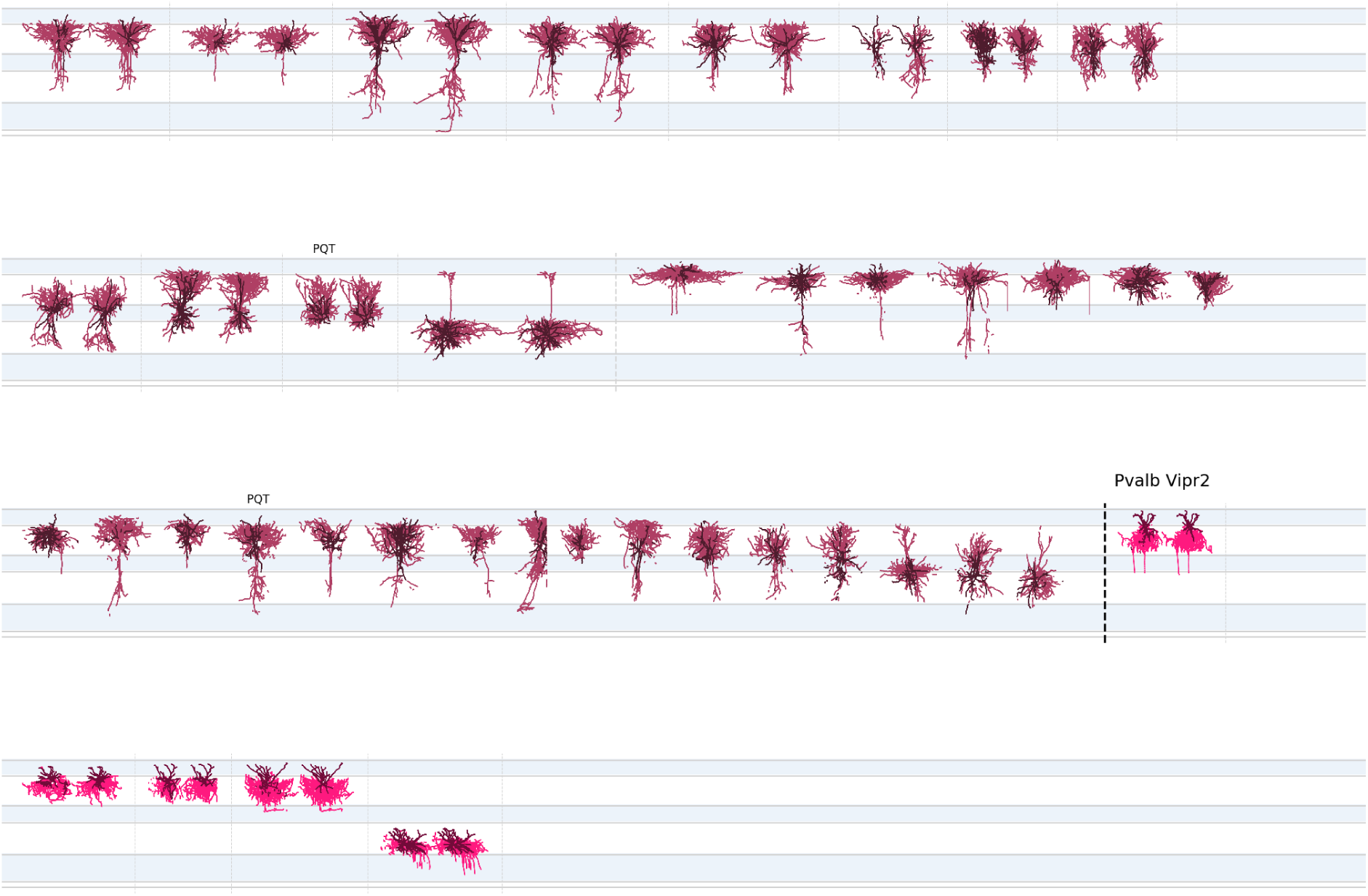
Morphological reconstructions of inhibitory neurons ordered by t-type – 11 of 11. Please refer to the caption of Figure S1.

**Figure S12:**
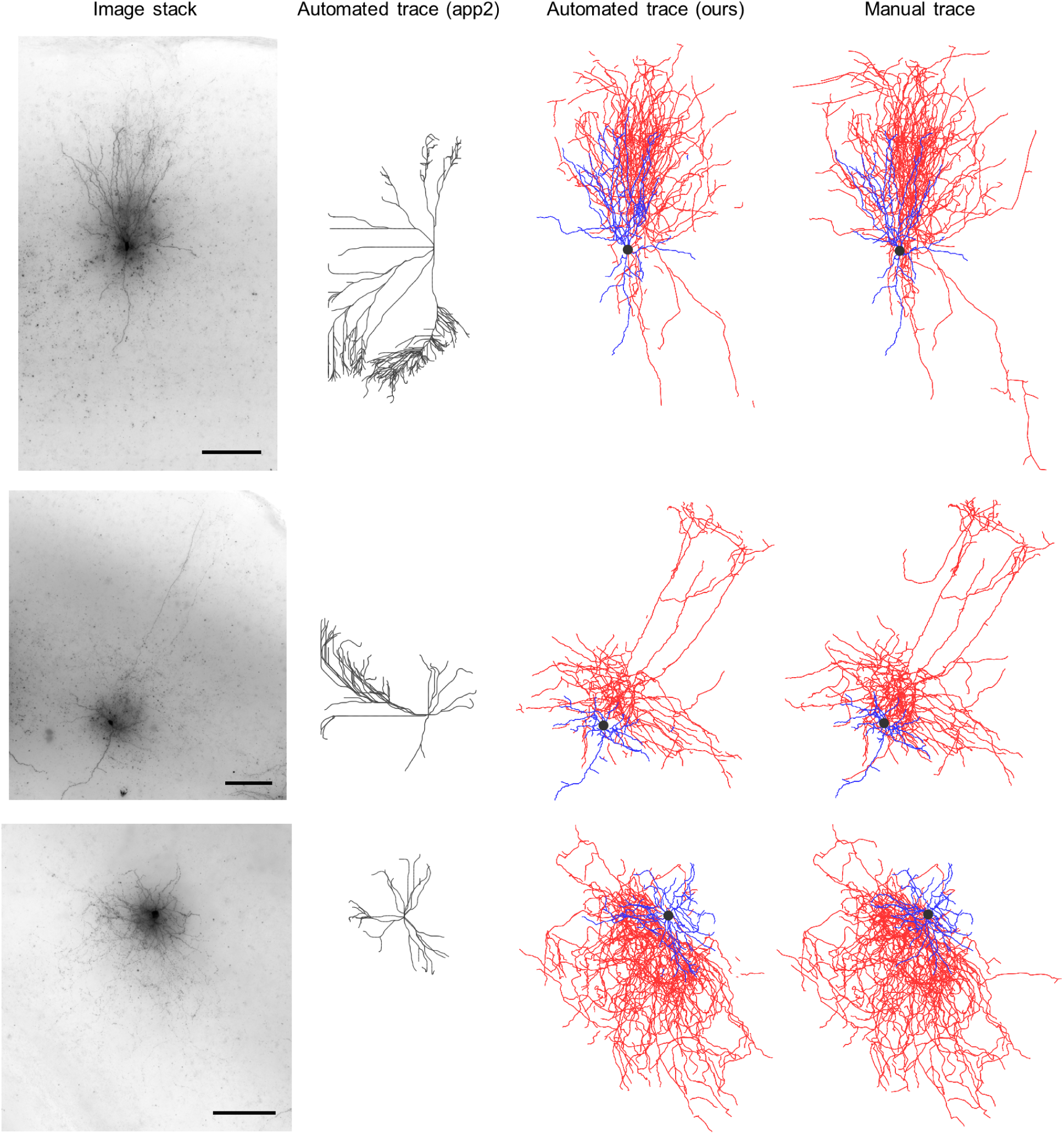
A qualitative comparative study of automated reconstruction accuracy based on three example test inhibitory neurons. Minimum intensity projection of image stacks (left column), automated reconstructions using the app2 algorithm^69, 70^ (second column), automated reconstructions using the proposed method (third column), manual reconstructions (right column). We tried multiple parameter values to optimize app2’s performance. For manual and our automated reconstructions: dendrites (blue), axons (red), soma (black). For automated reconstructions using app2: neurites (gray). Scale bar, 100 *µm*.

**Figure S13:**
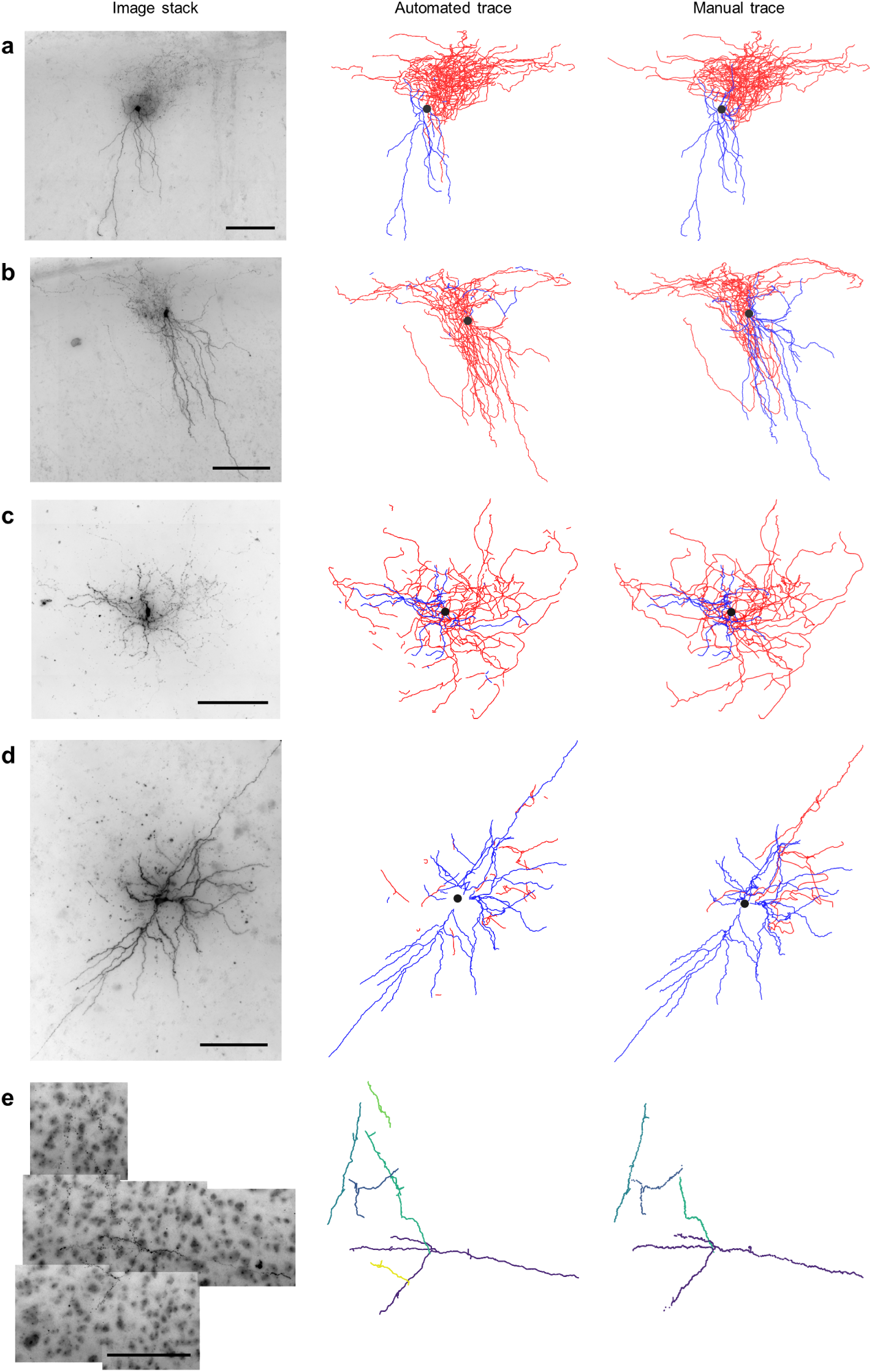
Morphological reconstructions of example neurons from other datasets, without tuning the segmentation model. Left column: minimum intensity projection of image stacks, middle column: automated reconstruction, right column: manual reconstruction. **a**, A human cortical inhibitory cell. **b**, Another human cortical inhibitory cell. All experimental steps prior to imaging were performed by a different laboratory and imaging was done at the Allen Institute^30^). **c**, A macaque subcortical inhibitory cell. **d**, Mouse subcortical inhibitory cell. Dendrites (blue), axons (red), soma (black). **e**, Part of a cat cortical excitatory cell, a part of the DIADEM dataset,^31^ originally obtained by Ref.^32^ – different colors indicate different connected components. Scale bar, 100 *µm*.

**Figure S14:**
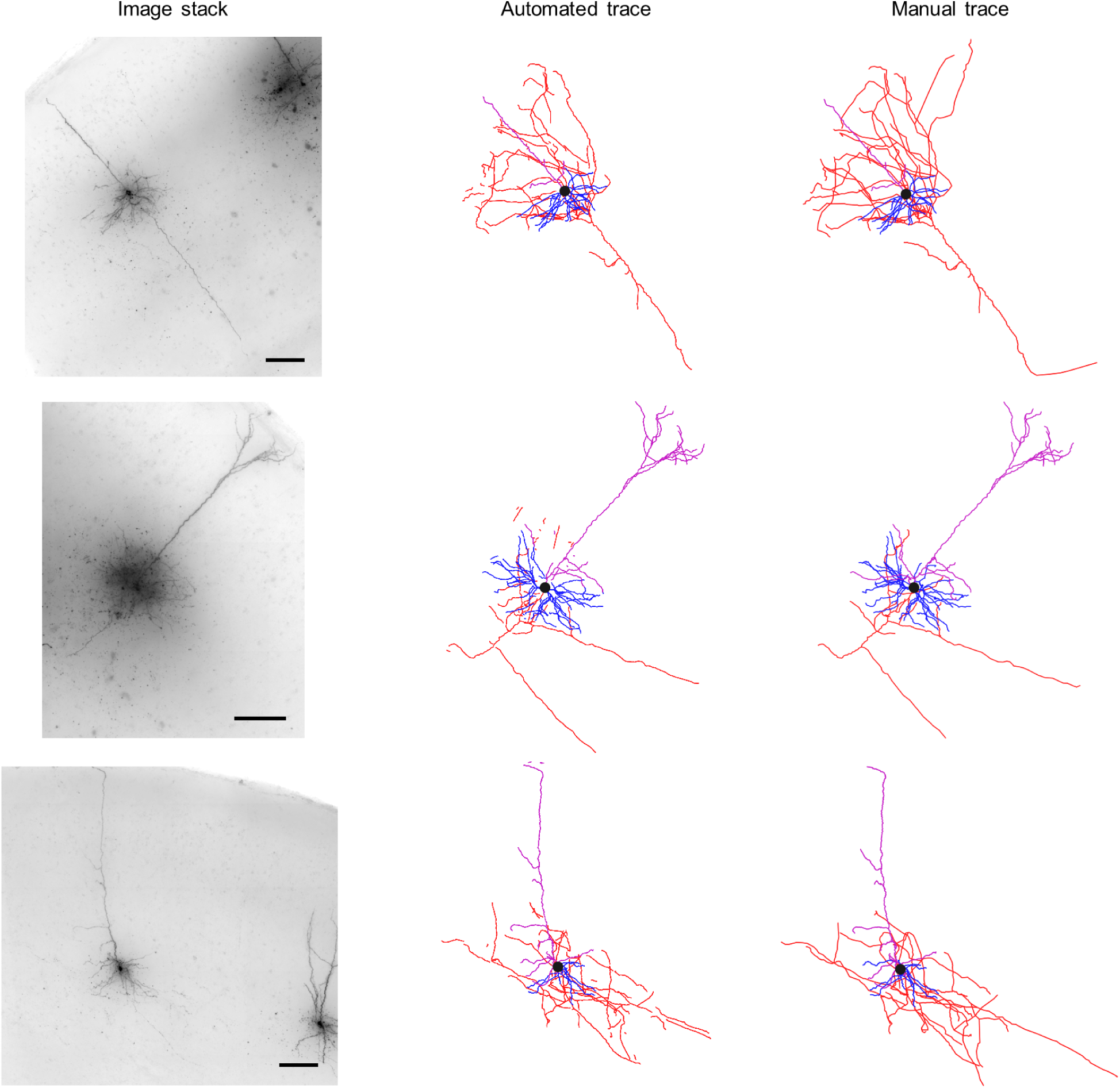
Morphological reconstructions of example excitatory neurons with traced axons. The method can trace and label the axon as well when it is captured in the slice. Minimum intensity projection of image stacks (left column), automated reconstructions (middle column), manual reconstructions (right column). Basal dendrites (blue), apical dendrites (magenta), axon (red), soma (black). Scale bar, 100 *µm*.

**Figure S15:**
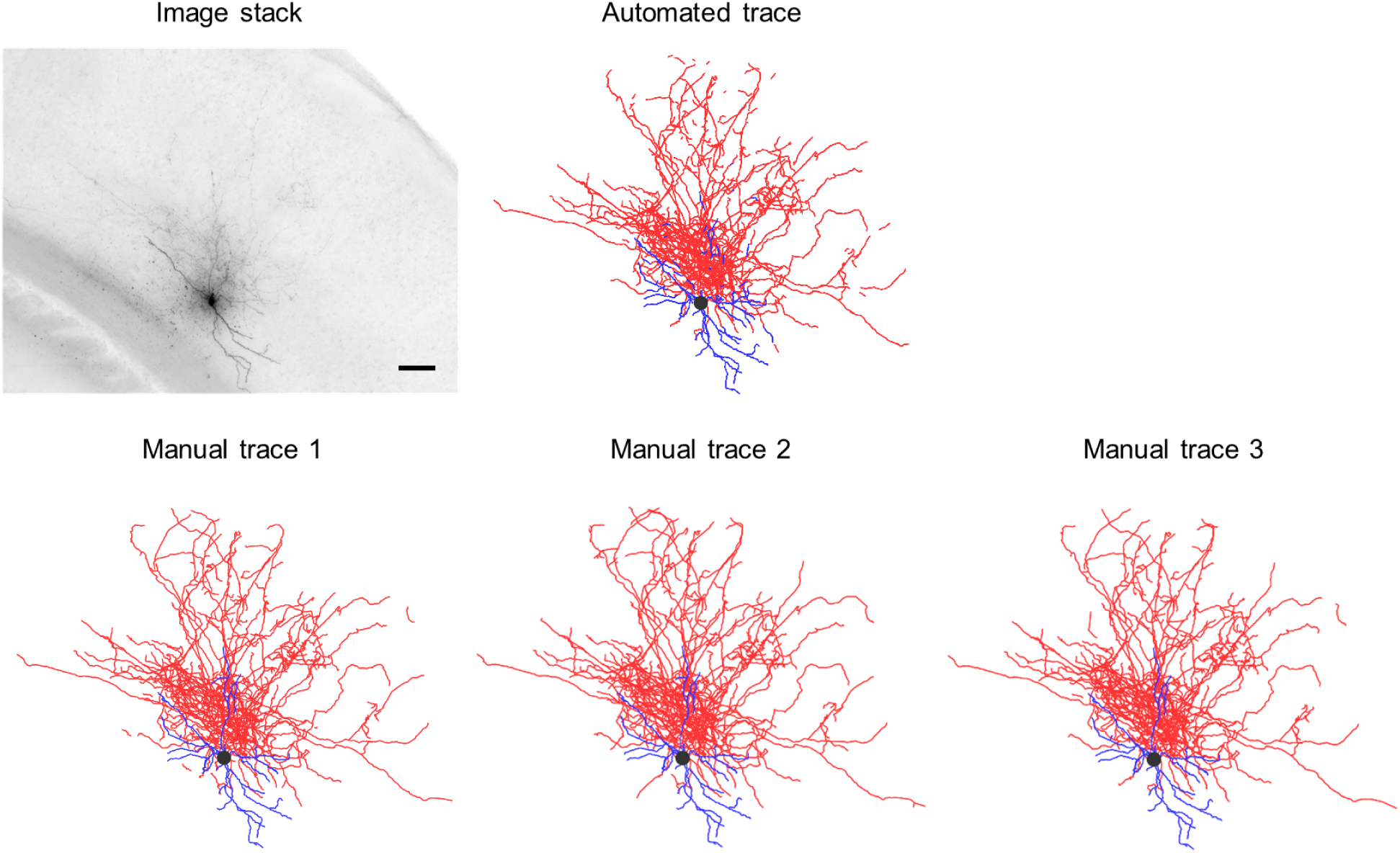
Comparison of automated vs. multiple manual reconstructions for one example test neuron. This neuron is not used for training. Multiple manual reconstructions are obtained to estimate cross-human agreement. Minimum intensity projection of the image stack (top left), automated reconstruction (top middle), manual reconstructions (bottom). Dendrites (blue), axons (red), soma (black). Scale bar, 100 *µm*. See Table S2 for quantification.

**Figure S16:**
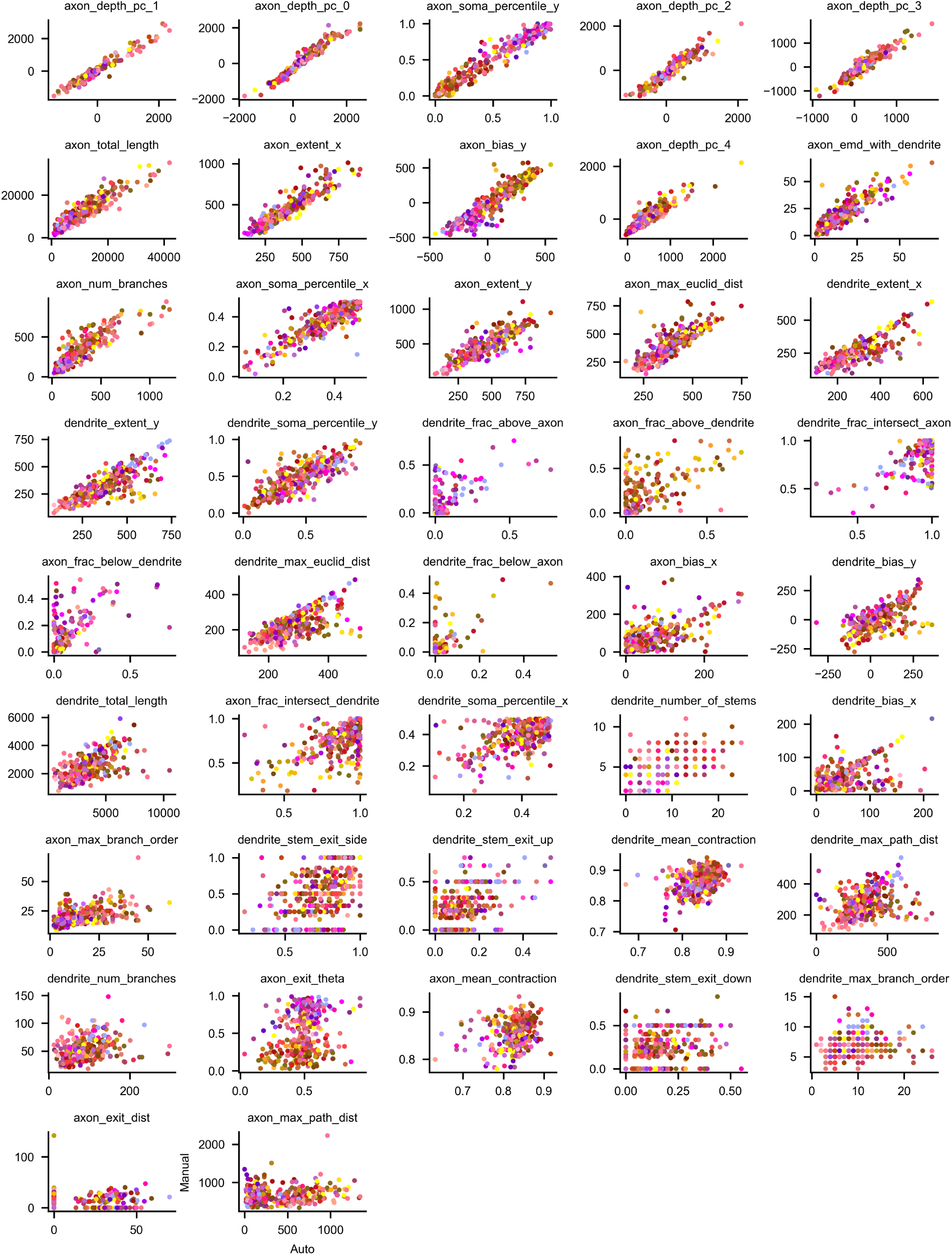
Comparison of automatically vs. manually generated morphometric features. Scatter plots for every feature are shown. For each feature, the *y*-axis denotes values based on manual traces and the *x*-axis denotes values based on automated traces. Each dot depicts an individual cell, and its color indicates the cell’s t-type, consistent with the coloring scheme in Ref.^5^

**Figure S17:**
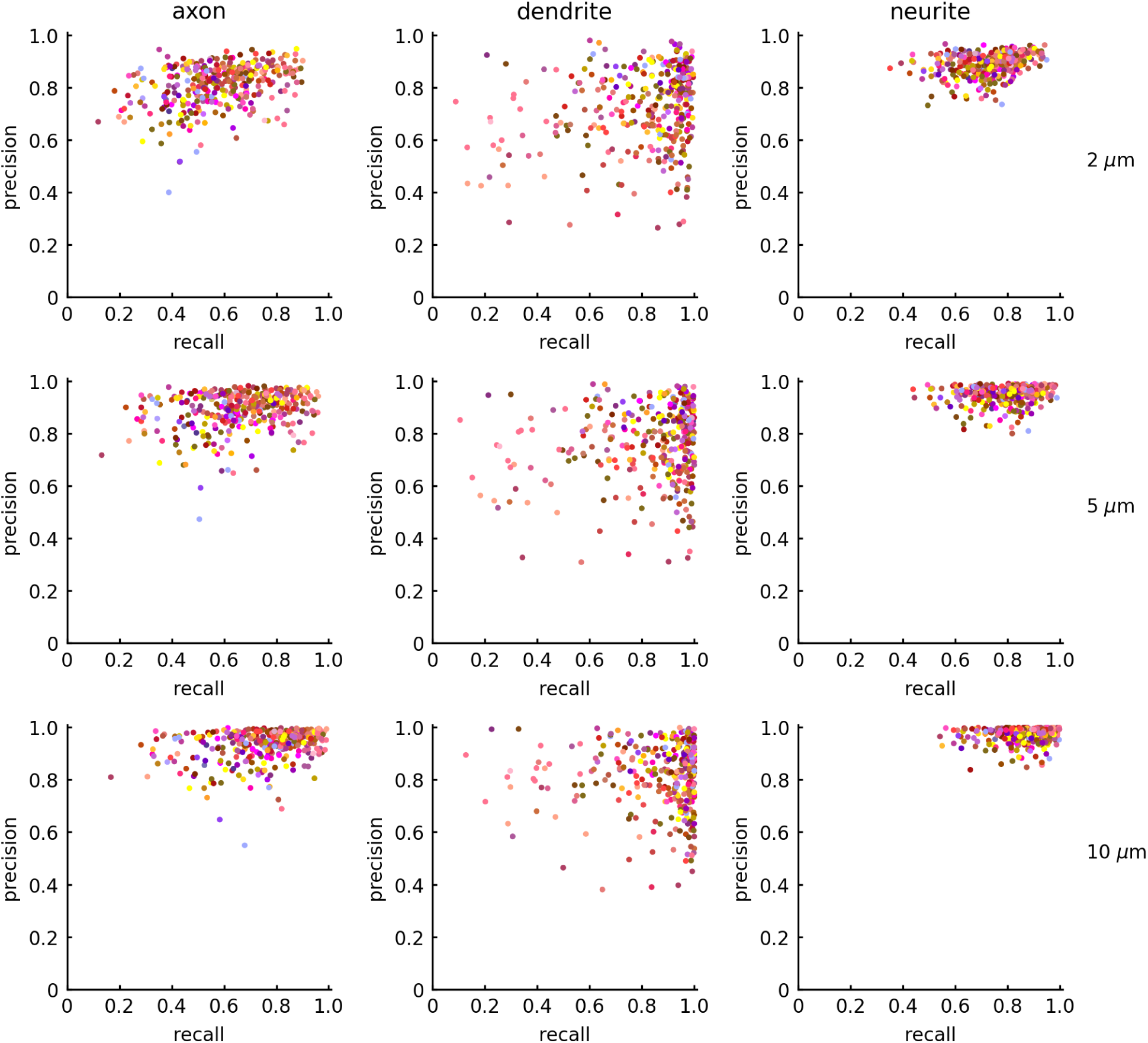
Neuron reconstruction accuracy. Scatter plots for precision vs. recall calculated by comparing automated and manual trace nodes within a given distance (2, 5, and 10 *µm*), as described in the main text, are shown for axonal, dendritic, and neurite (axonal or dendritic) nodes. Each dot depicts an individual cell, and its color indicates the cell’s t-type, consistent with the coloring scheme in Ref.^5^

**Figure S18:**
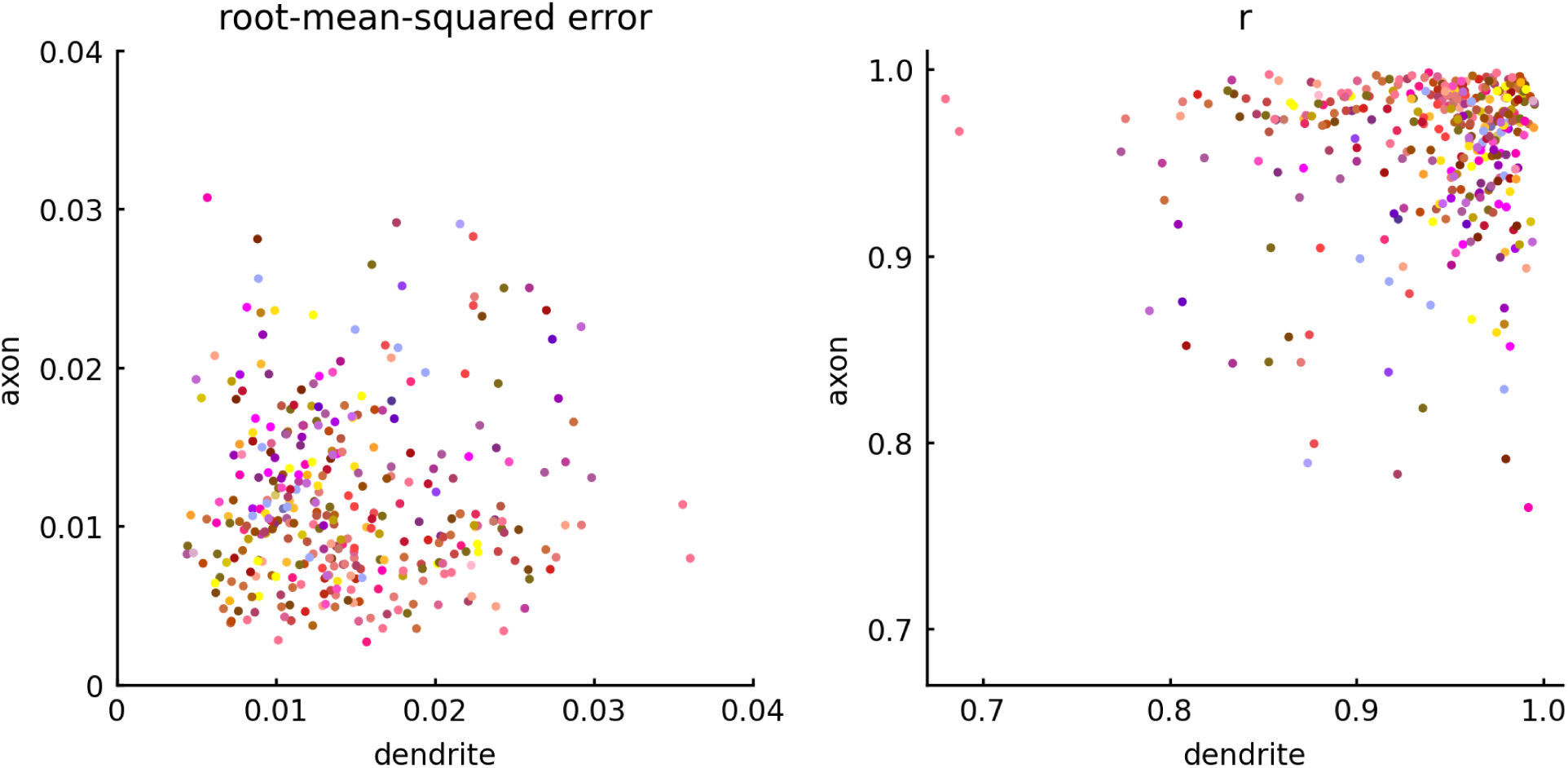
Comparison of automatically vs. manually generated ADRs. Scatter plots for the root-mean-squared error (left) and Pearson’s correlation *r* values (right) for each cell’s automatically reconstructed axonal vs. dendritic arbors, with respect to the corresponding manual traces, are shown. Each dot depicts an individual cell, and its color indicates the cell’s t-type, consistent with the coloring scheme in Ref.^5^

**Figure S19:**
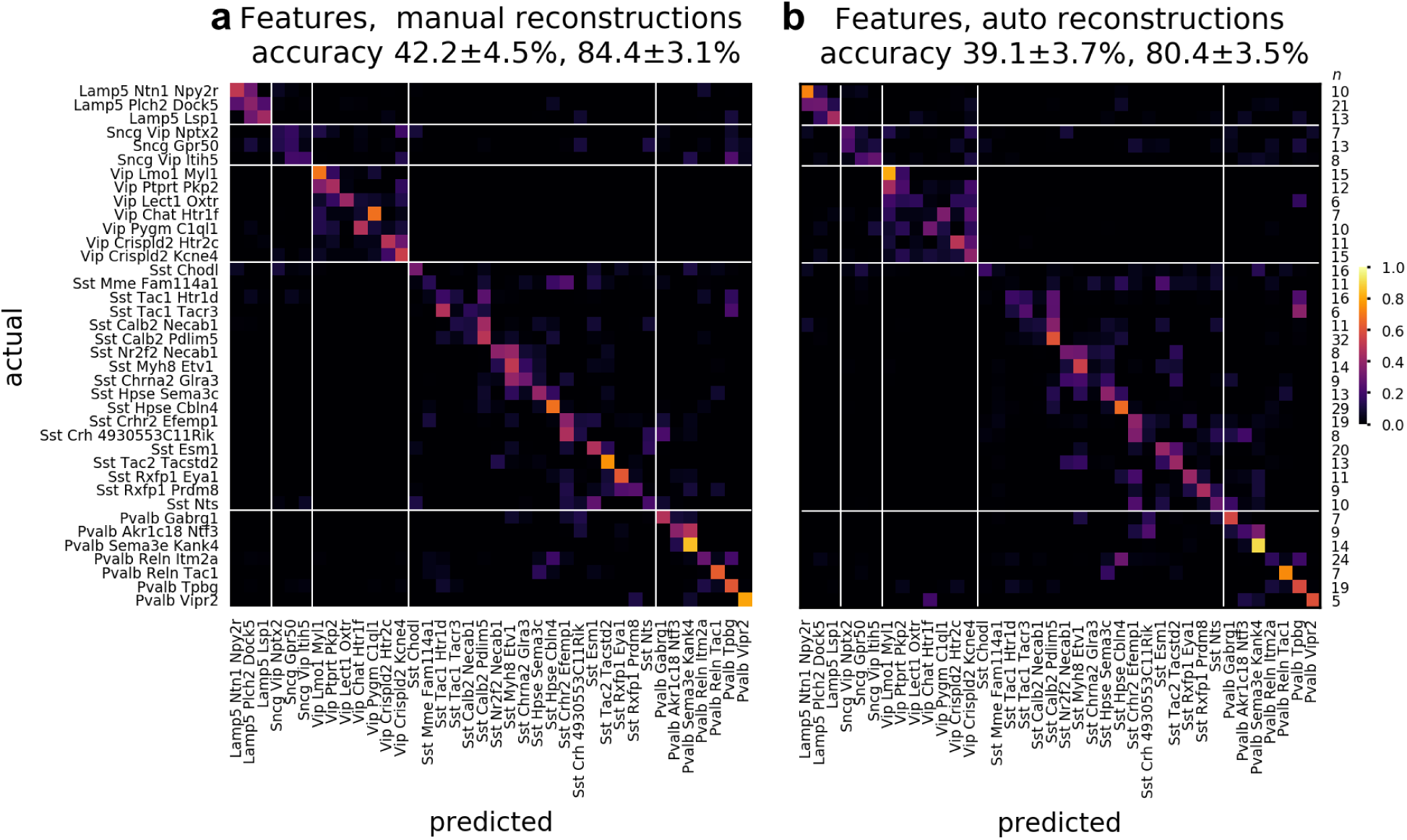
Comparison of cell type classification accuracy based on a set of classical morphometric features using manually vs. automatically reconstructed cells. Confusion matrix for the classification of 38 t-types based on features, using 488 manually (**a**) and automatically (**b**) reconstructed cells. Accuracy values reported in the headers refer to mean *±* s.d. of the overall t-type and t-subclass classifiers, respectively, across cross-validation folds. Rightmost column lists the number of cells in each t-type (*n*).

**Figure S20:**
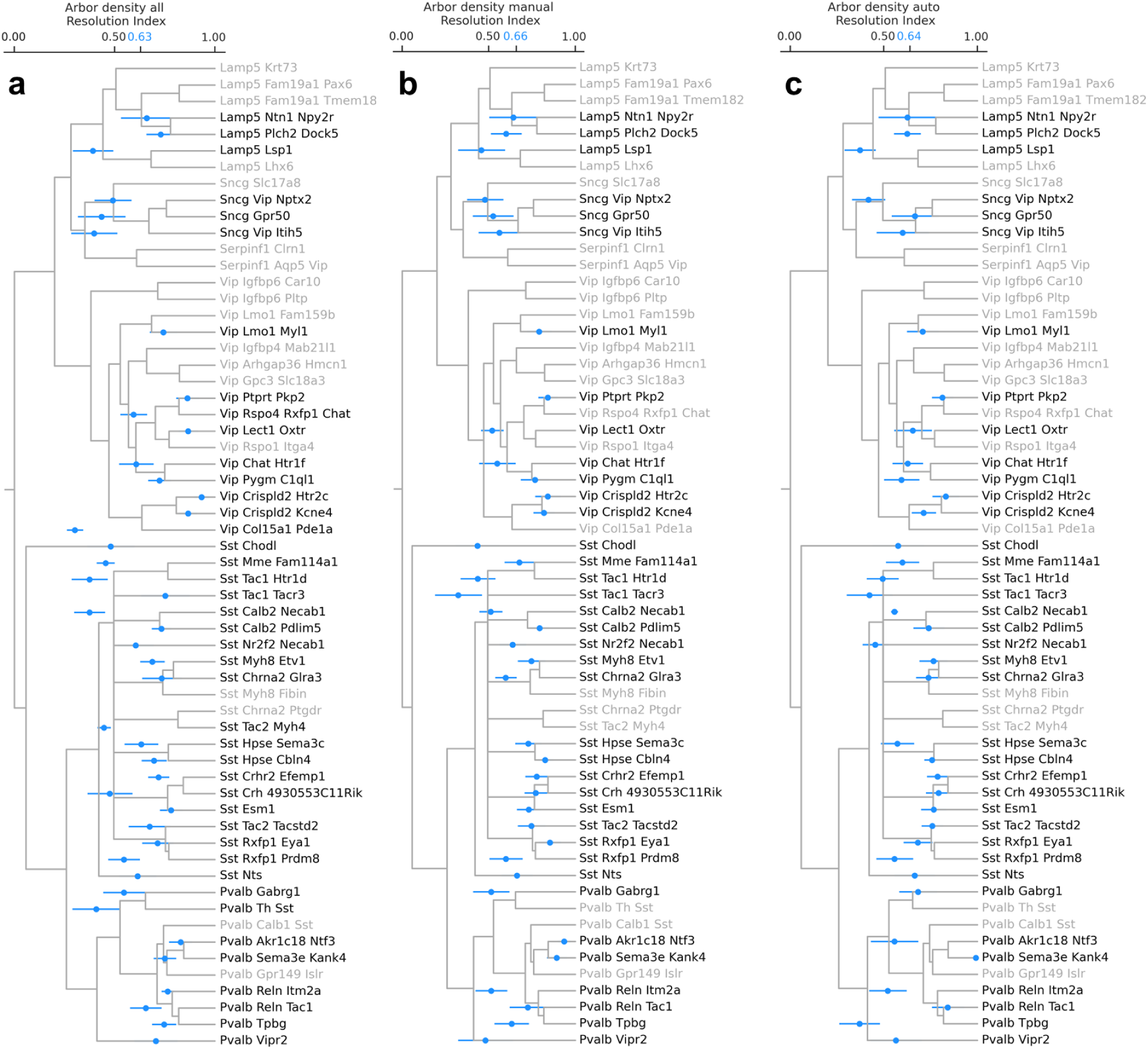
Arbor density-based cell type classification performance using resolution index. The depth of the Tasic et al. GABAergic neuron hierarchical tree is normalized to the 0-1 range, and resolution index for each cell is defined as the value of the closest common ancestor of the true and predicted leaf node labels. Perfect classification corresponds to resolution index of 1. Error bars indicate resolution index (mean *±* SE over 10-fold stratified cross validation sets) for each cell type. The overall mean across cell types is indicated by blue on the y-axis. Cell types absent from the dataset are indicated by grayed out labels on the x-axis. Classification was performed with multi-layer perceptron classifiers using arbor density representation of all (**a**), manually (**b**) and automatically (**c**) reconstructed neurons as input. Resolution index values are not strictly comparable across panels because the numbers of cells and types are not identical. Left panel: 747 cells, 42 types; middle and right panels: 488 cells, 38 types. (See Figure 3.)

**Figure S21:**
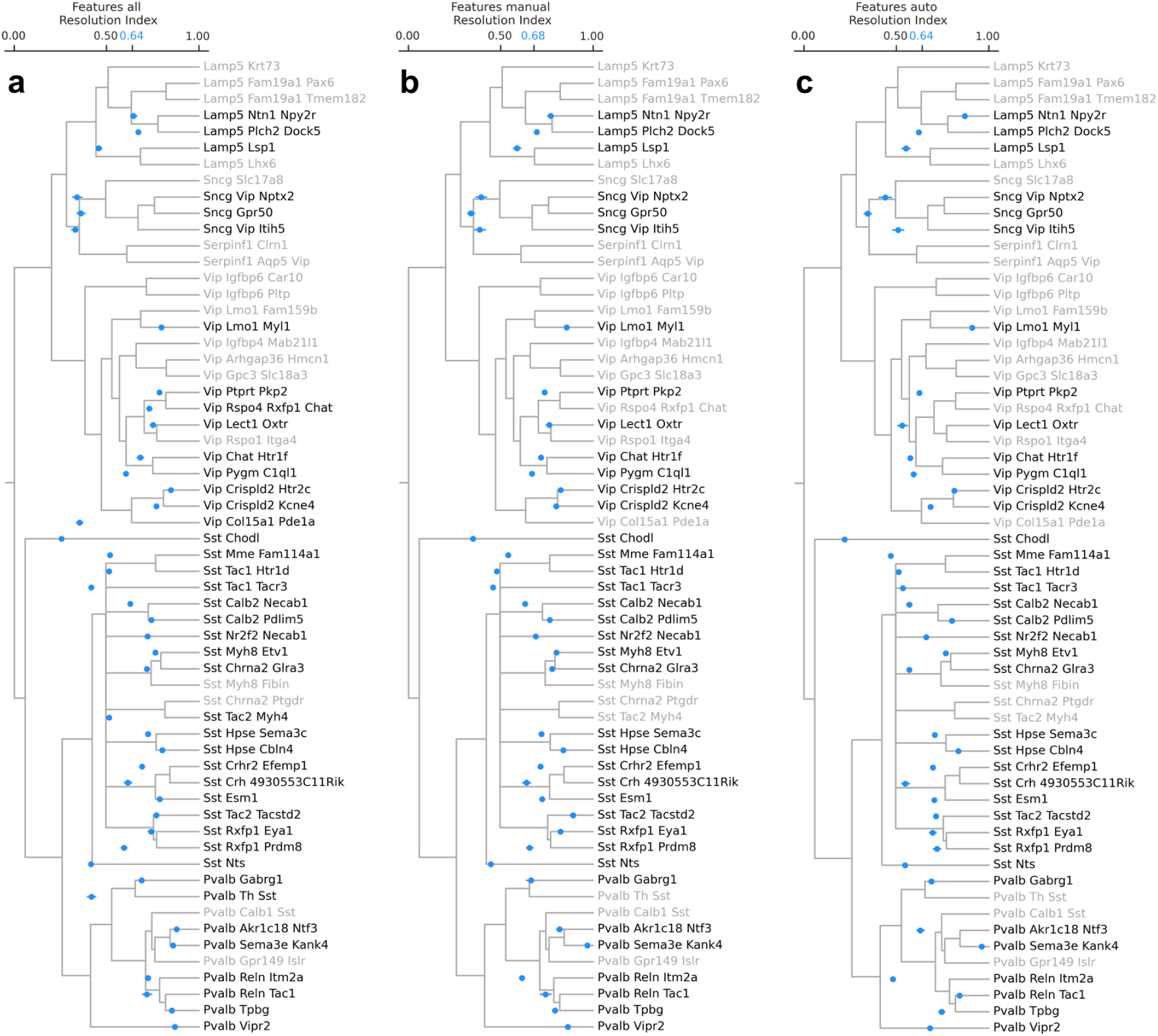
Feature-based cell type classification performance using resolution index. As before, error bars indicate resolution index (mean *±* SE over 5-fold stratified cross validation sets repeated 20 times) for each cell type. The overall mean across cell types is indicated by blue on the y-axis. Cell types absent from the dataset are indicated by grayed out labels on the x-axis. Classification was performed with random forest classifiers using morphometric features of all (**a**), manually (**b**) and automatically reconstructed cells (**c**) as input. Resolution index values are not strictly comparable across panels because the numbers of cells and types are not identical. Left panel: 747 cells, 42 types; middle and right panels: 488 cells, 38 types. (See Figure 3.)

**Table S1:** Neuron reconstruction accuracy. Precision, recall, and F1-score values are calculated by comparing automated and manual trace nodes within a given distance (2, 5, and 10 *µm*). Mean *±* s.d. of the values over 340 cells are reported for axonal, dendritic, and neurite (combined axonal and dendritic) nodes of raw and masked automated traces.

| **Precision** | Axon |  | Dendrite |  | Neurite |  |
| --- | --- | --- | --- | --- | --- | --- |
| Distance, $\mu m$ | raw | masked | raw | masked | raw | masked |
| 2 | $0.81 \pm 0.08$ | $0.83 \pm 0.07$ | $0.72 \pm 0.15$ | $0.73 \pm 0.15$ | $0.89 \pm 0.04$ | $0.91 \pm 0.03$ |
| 5 | $0.89 \pm 0.07$ | $0.91 \pm 0.06$ | $0.76 \pm 0.14$ | $0.77 \pm 0.14$ | $0.95 \pm 0.03$ | $0.97 \pm 0.02$ |
| 10 | $0.93 \pm 0.06$ | $0.96 \pm 0.05$ | $0.82 \pm 0.12$ | $0.82 \pm 0.12$ | $0.97 \pm 0.03$ | $0.99 \pm 0.01$ |

| **Recall** | Axon |  | Dendrite |  | Neurite |  |
| --- | --- | --- | --- | --- | --- | --- |
| Distance, $\mu m$ | raw | masked | raw | masked | raw | masked |
| 2 | $0.58 \pm 0.16$ | $0.58 \pm 0.16$ | $0.81 \pm 0.20$ | $0.81 \pm 0.20$ | $0.73 \pm 0.11$ | $0.73 \pm 0.11$ |
| 5 | $0.66 \pm 0.16$ | $0.66 \pm 0.16$ | $0.83 \pm 0.19$ | $0.83 \pm 0.19$ | $0.80 \pm 0.11$ | $0.80 \pm 0.11$ |
| 10 | $0.75 \pm 0.16$ | $0.75 \pm 0.16$ | $0.87 \pm 0.17$ | $0.87 \pm 0.17$ | $0.87 \pm 0.09$ | $0.87 \pm 0.09$ |

Table S1: **Neuron reconstruction accuracy.**
| **F1-score** | Axon |  | Dendrite |  | Neurite |  |
| --- | --- | --- | --- | --- | --- | --- |
| Distance, $\mu m$ | raw | masked | raw | masked | raw | masked |
| 2 | $0.66 \pm 0.13$ | $0.67 \pm 0.13$ | $0.74 \pm 0.16$ | $0.74 \pm 0.16$ | $0.80 \pm 0.08$ | $0.80 \pm 0.08$ |
| 5 | $0.75 \pm 0.13$ | $0.76 \pm 0.13$ | $0.78 \pm 0.15$ | $0.78 \pm 0.15$ | $0.87 \pm 0.07$ | $0.87 \pm 0.07$ |
| 10 | $0.82 \pm 0.12$ | $0.83 \pm 0.12$ | $0.83 \pm 0.13$ | $0.83 \pm 0.13$ | $0.92 \pm 0.06$ | $0.92 \pm 0.06$ |

**Table S2:** Neuron reconstruction accuracy for multiple manual traces vs. the automated trace for a single test cell. See Figure S15 for projections of the raw image and the corresponding reconstructions for this test neuron. Precision, recall, and F1-score values are calculated by comparing pairs of trace nodes within a given distance (2, 5, and 10 *µm*). Mean *±* s.d. of the values over three pairwise comparisons are reported for axonal, dendritic, and neurite (combined axonal and dendritic) nodes of manual vs. manual traces and automated vs. manual traces.

| **Precision** | Axon |  | Dendrite |  | Neurite |  |
| --- | --- | --- | --- | --- | --- | --- |
| Distance, $\mu m$ | manual | auto | manual | auto | manual | auto |
| 2 | $0.93 \pm 0.02$ | $0.88 \pm 0.01$ | $0.99 \pm 0.00$ | $0.82 \pm 0.00$ | $0.93 \pm 0.02$ | $0.91 \pm 0.01$ |
| 5 | $0.97 \pm 0.01$ | $0.92 \pm 0.01$ | $1.00 \pm 0.00$ | $0.86 \pm 0.01$ | $0.97 \pm 0.01$ | $0.94 \pm 0.01$ |
| 10 | $0.98 \pm 0.01$ | $0.95 \pm 0.01$ | $1.00 \pm 0.00$ | $0.89 \pm 0.01$ | $0.98 \pm 0.01$ | $0.96 \pm 0.01$ |

| **Recall** | Axon |  | Dendrite |  | Neurite |  |
| --- | --- | --- | --- | --- | --- | --- |
| Distance, $\mu m$ | manual | auto | manual | auto | manual | auto |
| 2 | $0.92 \pm 0.01$ | $0.91 \pm 0.01$ | $0.98 \pm 0.00$ | $0.65 \pm 0.00$ | $0.92 \pm 0.01$ | $0.92 \pm 0.01$ |
| 5 | $0.96 \pm 0.02$ | $0.96 \pm 0.01$ | $0.99 \pm 0.00$ | $0.71 \pm 0.00$ | $0.96 \pm 0.01$ | $0.98 \pm 0.01$ |
| 10 | $0.97 \pm 0.01$ | $0.99 \pm 0.00$ | $0.99 \pm 0.00$ | $0.73 \pm 0.00$ | $0.97 \pm 0.01$ | $0.99 \pm 0.00$ |

Table S2: **Neuron reconstruction accuracy for multiple manual traces vs.**
| **F1-score** | Axon |  | Dendrite |  | Neurite |  |
| --- | --- | --- | --- | --- | --- | --- |
| Distance, $\mu m$ | manual | auto | manual | auto | manual | auto |
| 2 | $0.92 \pm 0.01$ | $0.89 \pm 0.00$ | $0.99 \pm 0.00$ | $0.73 \pm 0.00$ | $0.93 \pm 0.01$ | $0.91 \pm 0.00$ |
| 5 | $0.96 \pm 0.00$ | $0.94 \pm 0.00$ | $0.99 \pm 0.00$ | $0.78 \pm 0.00$ | $0.97 \pm 0.00$ | $0.96 \pm 0.00$ |
| 10 | $0.98 \pm 0.00$ | $0.97 \pm 0.00$ | $1.00 \pm 0.00$ | $0.80 \pm 0.00$ | $0.98 \pm 0.00$ | $0.98 \pm 0.00$ |

**Table S3:** Statistical significance and effect size values for predicting anatomical features from gene expression via sparse linear regression for seven different cell types and four subclasses. For each entry, the FDR-corrected *p*-value as calculated by a non-parametric shuffle test is listed. (See Sparse feature selection analysis under Methods.) If the value is considered statistically significant at *p ≤* 0.05, the *R*^2^ value is also displayed (*p* / *R*^2^). *p*-values less than or equal to 0.05 and *R*^2^ values larger than or equal to 0.25 are shown in bold.

|  | Sst Calb2 Pdlm5 | Sst Hpse Cbln4 | Sst Chodl | Pvalb Reln Itm2a | Pvalb Tpbq | Lamp5 Lsp1 |
| --- | --- | --- | --- | --- | --- | --- |
| L1 axon | **0 / 0.40** | 1.000 | 0.637 | **0.023 / 0.13** | **0.033 / 0.21** | **0.013 / 0.56** |
| L2/3 axon | **0.013 / 0.30** | 1.000 | 1.000 | **0 / 0.35** | **0.023 / 0.27** | **0.042 / 0.52** |
| L4 axon | 0.689 | 0.332 | 0.607 | **0 / 0.35** | 0.216 | 1.000 |
| L5 axon | **0.033 / 0.15** | **0.013 / 0.25** | 1.000 | 0.102 | 0.058 | 0.135 |
| L6 axon | 1.000 | 0.081 | 0.288 | 0.182 | 1.000 | 0.058 |
| L1 dendrite | **0.042 / 0.17** | 1.000 | 0.689 | 0.088 | **0 / 0.42** | 0.283 |
| L2/3 dendrite | **0.033 / 0.15** | 1.000 | 0.332 | **0.013 / 0.27** | **0.042 / 0.21** | 1.000 |
| L4 dendrite | 0.697 | **0 / 0.40** | **0.013 / 0.40** | 0.058 | 0.074 | 0.393 |
| L5 dendrite | **0 / 0.19** | 0.102 | 0.393 | **0 / 0.25** | **0.013 / 0.26** | 1.000 |
| L6 dendrite | 1.000 | 0.088 | 0.866 | 0.051 | 0.363 | 1.000 |
| soma depth | **0 / 0.26** | **0.023 / 0.21** | 0.246 | **0 / 0.34** | **0.013 / 0.29** | **0.023 / 0.48** |
| axon centroid | **0.023 / 0.16** | 0.210 | 0.058 | **0 / 0.34** | 0.058 | **0.013 / 0.44** |
| dendrite centroid | 0.150 | **0.023 / 0.26** | 0.096 | **0 / 0.39** | **0.013 / 0.38** | **0.023 / 0.46** |
|  | Lamp5 Plch2 Dock5 | Sst | Pvalb | Lamp5 | Vip |  |
| L1 axon | **0.042 / 0.40** | **0 / 0.29** | **0 / 0.29** | **0 / 0.47** | **0 / 0.24** |  |
| L2/3 axon | 0.322 | **0 / 0.41** | **0 / 0.66** | 0.813 | **0 / 0.27** |  |
| L4 axon | 0.351 | **0 / 0.44** | **0 / 0.40** | 0.210 | **0 / 0.18** |  |
| L5 axon | 0.081 | **0 / 0.33** | **0 / 0.21** | 1.000 | **0 / 0.18** |  |
| L6 axon | 0.540 | **0 / 0.37** | **0 / 0.52** | 1.000 | 0.813 |  |
| L1 dendrite | **0 / 0.40** | **0.023 / 0.01** | **0 / 0.42** | **0 / 0.25** | 1.000 |  |
| L2/3 dendrite | 1.000 | **0 / 0.37** | **0 / 0.66** | 1.000 | **0 / 0.48** |  |
| L4 dendrite | 1.000 | **0 / 0.42** | **0 / 0.28** | 1.000 | **0 / 0.16** |  |
| L5 dendrite | 1.000 | **0 / 0.19** | **0 / 0.24** | 1.000 | **0 / 0.34** |  |
| L6 dendrite | 1.000 | **0 / 0.44** | **0 / 0.59** | 1.000 | 0.689 |  |
| soma depth | 0.067 | **0 / 0.48** | **0 / 0.70** | **0 / 0.24** | **0 / 0.33** |  |
| axon centroid | 0.074 | **0 / 0.53** | **0 / 0.72** | **0 / 0.25** | **0 / 0.29** |  |
| dendrite centroid | 0.074 | **0 / 0.45** | **0 / 0.68** | **0 / 0.26** | **0 / 0.21** |  |

**Table S4:** Statistical significance and effect size values for predicting anatomical features from gene expression via sparse linear regression for seven different cell types and four subclasses – before FDR correction. For each entry, original *p*-value as calculated by a non-parametric shuffle test is listed. If the value is considered statistically significant at *p ≤* 0.05, the *R*^2^ value is also displayed (*p* / *R*^2^). *p*-values less than or equal to 0.05 and *R*^2^ values larger than or equal to 0.25 are shown in bold.

|  | Sst Calb2 Pdlm5 | Sst Hpse Cbln4 | Sst Chodl | Pvalb Reln Itm2a | Pvalb Tpbgr | Lamp5 Lsp1 |
| --- | --- | --- | --- | --- | --- | --- |
| L1 axon | **0 / 0.40** | 0.481 | 0.090 | **0.002 / 0.13** | **0.003 / 0.21** | **0.001 / 0.56** |
| L2/3 axon | **0.001 / 0.30** | 0.232 | 0.306 | **0 / 0.35** | **0.002 / 0.27** | **0.004 / 0.52** |
| L4 axon | 0.100 | **0.044 / 0.01** | 0.085 | **0 / 0.35** | **0.027 / 0.06** | 0.312 |
| L5 axon | **0.003 / 0.15** | **0.001 / 0.25** | 0.347 | **0.012 / 0.07** | **0.006 / 0.14** | **0.016 / -0.45** |
| L6 axon | 0.251 | **0.009 / 0.10** | **0.037 / 0.11** | **0.022 / -0.05** | 0.227 | **0.006 / 0.32** |
| L1 dendrite | **0.004 / 0.17** | 0.895 | 0.100 | **0.010 / -0.26** | **0 / 0.42** | **0.036 / -0.03** |
| L2/3 dendrite | **0.003 / 0.15** | 0.234 | **0.044 / 0.05** | **0.001 / 0.27** | **0.004 / 0.21** | 0.189 |
| L4 dendrite | 0.102 | **0 / 0.40** | **0.001 / 0.40** | **0.006 / 0.09** | **0.008 / 0.15** | 0.054 |
| L5 dendrite | **0 / 0.19** | **0.012 / 0.13** | 0.054 | **0 / 0.25** | **0.001 / 0.26** | 0.174 |
| L6 dendrite | 0.230 | **0.010 / 0.15** | 0.130 | **0.005 / 0.08** | **0.049 / 0.01** | 0.177 |
| soma depth | **0 / 0.26** | **0.002 / 0.21** | **0.031 / 0.06** | **0 / 0.34** | **0.001 / 0.29** | **0.002 / 0.48** |
| axon centroid | **0.002 / 0.16** | **0.026 / 0.04** | **0.006 / 0.26** | **0 / 0.34** | **0.006 / 0.21** | **0.001 / 0.44** |
| dendrite centroid | **0.018 / 0.18** | **0.002 / 0.26** | **0.011 / 0.22** | **0 / 0.39** | **0.001 / 0.38** | **0.002 / 0.46** |
|  | Lamp5 Plch2 Dock5 | Sst | Pvalb | Lamp5 | Vip |  |
| L1 axon | **0.004 / 0.40** | **0 / 0.29** | **0 / 0.29** | **0 / 0.47** | **0 / 0.24** |  |
| L2/3 axon | **0.042 / 0.08** | **0 / 0.41** | **0 / 0.66** | 0.120 | **0 / 0.27** |  |
| L4 axon | **0.047 / 0.01** | **0 / 0.44** | **0 / 0.40** | **0.026 / 0.01** | **0 / 0.18** |  |
| L5 axon | **0.009 / 0.07** | **0 / 0.33** | **0 / 0.21** | 0.186 | **0 / 0.18** |  |
| L6 axon | 0.075 | **0 / 0.37** | **0 / 0.52** | 0.177 | 0.121 |  |
| L1 dendrite | **0 / 0.40** | **0.002 / 0.01** | **0 / 0.42** | **0 / 0.25** | 0.208 |  |
| L2/3 dendrite | 0.218 | **0 / 0.37** | **0 / 0.66** | 0.510 | **0 / 0.48** |  |
| L4 dendrite | 0.285 | **0 / 0.42** | **0 / 0.28** | 0.214 | **0 / 0.16** |  |
| L5 dendrite | 0.217 | **0 / 0.19** | **0 / 0.24** | 0.217 | **0 / 0.34** |  |
| L6 dendrite | 0.292 | **0 / 0.44** | **0 / 0.59** | 0.160 | 0.099 |  |
| soma depth | **0.007 / 0.20** | **0 / 0.48** | **0 / 0.70** | **0 / 0.24** | **0 / 0.33** |  |
| axon centroid | **0.008 / 0.23** | **0 / 0.53** | **0 / 0.72** | **0 / 0.25** | **0 / 0.29** |  |
| dendrite centroid | **0.008 / 0.16** | **0 / 0.45** | **0 / 0.68** | **0 / 0.26** | **0 / 0.21** |  |

**Table S5:**
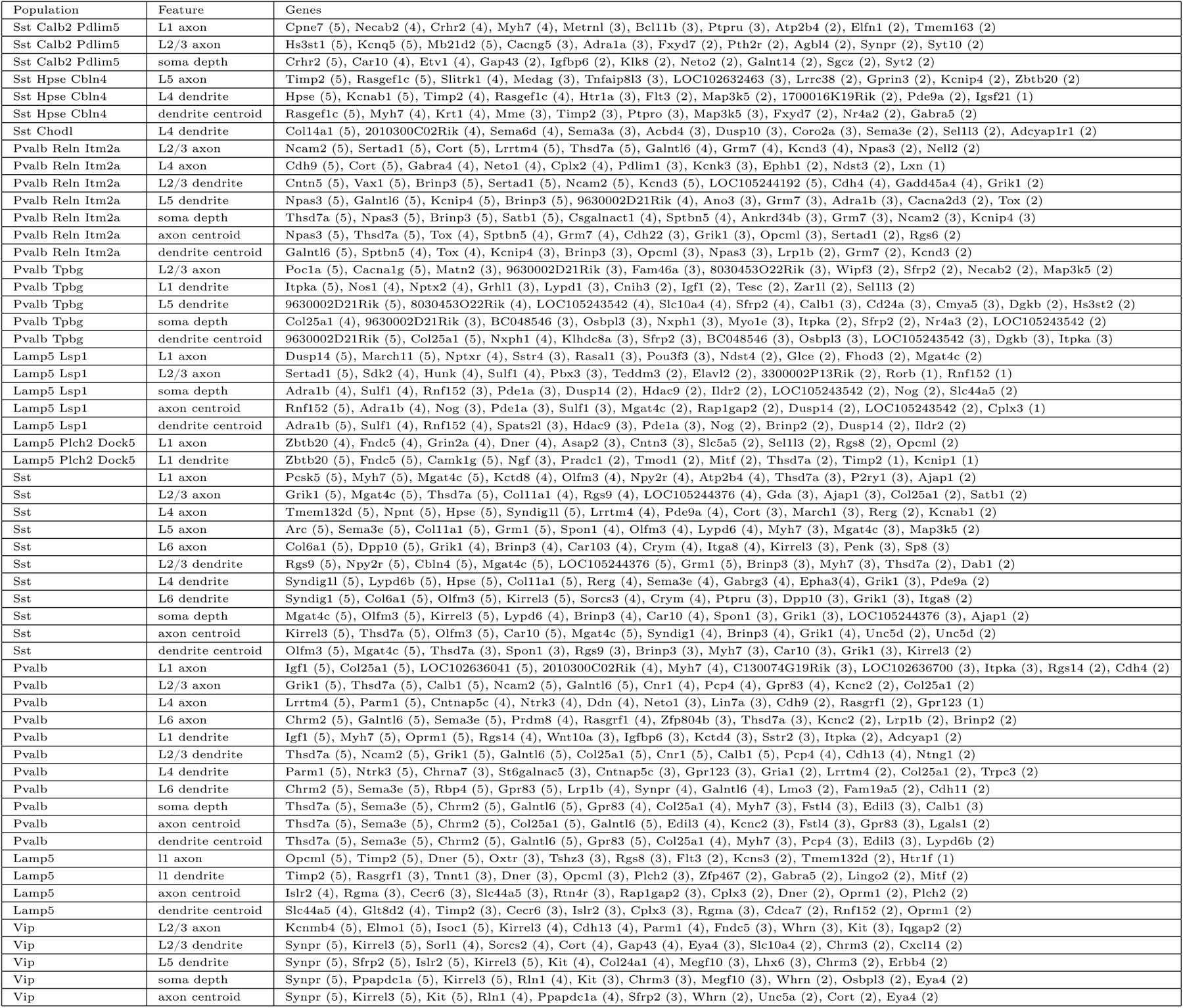
Gene sets selected via sparse linear regression for different cell types/subclasses and anatomical features. Only statistically significant sets with R^2^ _ 0:25 are shown (See Supplementary Table 1). Numbers in parentheses denote the number of times the preceding gene was selected out of 5 cross-validation runs.

